# Notch-dependent left-right asymmetry in β-catenin-TCF activity instructs anteroposterior positioning of motor neurons in the *C. elegans* ventral nerve cord

**DOI:** 10.1101/2025.03.02.641079

**Authors:** Wesley Chan, Justin Evans, Saber Saharkhiz, Clover Stubbert, Elicia Preston, Jonathan D. Rumley, Tony Roenspies, Pavak K. Shah, John I. Murray, Theodore J. Perkins, Antonio Colavita

## Abstract

The precise organization of motor neurons along the neuroaxis is essential for nervous system connectivity and function. In newly hatched *C. elegans*, the ventral nerve cord (VNC) contains 22 motor neurons of three classes, DD, DA, and DB, whose cell bodies are arranged in a stereotyped pattern along the anteroposterior (AP) axis. However, the mechanisms that control motor neuron positioning during nerve cord assembly are not fully understood. Here, we uncover a left-right asymmetry in β-catenin-TCF transcriptional activity that regulates AP motor neuron positioning. Loss of *bar-1*/β-catenin or *pop-1*/TCF causes a shift of motor neuron cell bodies toward the anterior, while loss of *pry-1*/Axin shifts them toward the posterior. We further find that POP-1/TCF activity is asymmetric between the left and right sides of the VNC in a BAR-1-dependent manner. Increasing BAR-1 levels on both sides of the VNC causes a posterior shift in motor neuron cell body placement. During VNC assembly, asymmetric Notch signaling promotes PRY-1 expression in left-side neurons, thereby restricting BAR-1 accumulation to right-side motor neurons, presumably through degradation by the destruction complex. Together, our findings reveal a previously unrecognized pathway in which an initial Notch-dependent left-right asymmetry establishes asymmetric β-catenin-TCF activity to regulate the AP positioning of motor neuron cell bodies along the VNC.

## INTRODUCTION

Central nerve cord development involves the organization of diverse motor neuron types along the neuroaxis to enable neuromuscular control along the full length of the body. An essential aspect of this process is ensuring the proper sorting and positioning of motor neuron cell bodies within each segment. Several mechanisms have been shown to contribute to neuronal positioning. Various attractive and repulsive guidance pathways, including stop signals, direct the migration of neurons to their final positions (Ding et al., 2005; Junge et al., 2016). Differential adhesive interactions sort neurons into specific pools or anchor them to specific locations (Bello et al., 2012; Demireva et al., 2011; Dewitz et al., 2018). Physical barriers, such as axon tracts or boundary cells, can further constrain cell body positioning (Bron et al., 2007; Laumonnerie et al., 2015; Lee and Song, 2013). The spatiotemporal expression of the molecules that mediate these activities is itself regulated by transcription factors that pattern the neuroaxis (Sagner and Briscoe, 2019), although the mechanisms that coordinate these positional cues during motor neuron assembly remain incompletely understood.

In *C. elegans*, although the ventral nerve cord (VNC) is not segmented, it contains a repeating organization of motor neurons along its length, which innervate ventral and dorsal muscle bands to generate sinusoidal-like movement (Lu et al., 2022). At hatching, the VNC contains 22 motor neurons belonging to three classes: 6 GABAergic DD, 9 cholinergic DA, and 7 cholinergic DB neurons (J E Sulston et al., 1983). The cell bodies of these neurons are positioned in a stereotypical manner (Lu et al., 2022; Saharkhiz et al., 2025). This arrangement is formed through a coordinated series of movements by developing neurons starting at the bean stage of embryonic development (Shah et al., 2017). At this stage, DD, DA and DB neurons born on the left and right sides form a dynamic monolayer, with central DB neurons flanked by DD and DA neurons on each side. These cells undergo convergent extension movements, intercalating towards the midline to assemble the presumptive VNC (Shah et al., 2017). By the 1.5-fold stage, the VNC is organized into two layers, with DD and DA neurons forming the dorsal layer and DB neurons forming the ventral layer (J. E. Sulston et al., 1983). DB neurons subsequently intercalate among the DD and DA neurons to assemble a single, stereotypically arranged tract of motor neurons. Perturbations during assembly are readily identifiable as changes in the position or arrangement of motor neuron cell bodies at hatching, providing a sensitive readout for genes that control VNC morphogenesis (Shah et al., 2017).

Here, we identify a pathway in which asymmetric Notch signaling establishes left-right (LR) asymmetry in β-catenin-TCF activity to regulate motor neuron positioning during VNC assembly. Genetic analysis of this pathway revealed that loss of *bar-1*/β-catenin or *pop-1*/TCF causes anterior shifts in motor neuron cell bodies, whereas loss of *pry-1*/Axin leads to posterior shifts. We find that asymmetric Notch signaling in left-side neurons, promotes expression of the destruction complex component PRY-1, thereby restricting BAR-1 accumulation to the right side of the developing VNC. This BAR-1 asymmetry is reflected in POP-1 transcriptional activity, which differs between the two sides of the VNC in a BAR-1-dependent manner. Increasing BAR-1 on both sides of the VNC causes posterior shifts in motor neuron positioning. Our findings identify a mechanism in which Notch signaling regulates BAR-1-POP-1 activity through asymmetric control of PRY-1 expression, distinct from the canonical Wnt pathway in which Wnt signaling regulates the destruction complex to control β-catenin stability. Ultimately, this asymmetry influences cell intercalation behaviours that establish the stereotyped organization of motor neurons along the AP axis. Together these findings show how LR asymmetry during neuroaxis formation can determine AP motor neuron positioning, revealing an unexpected link between LR and AP axis organization in the developing nervous system.

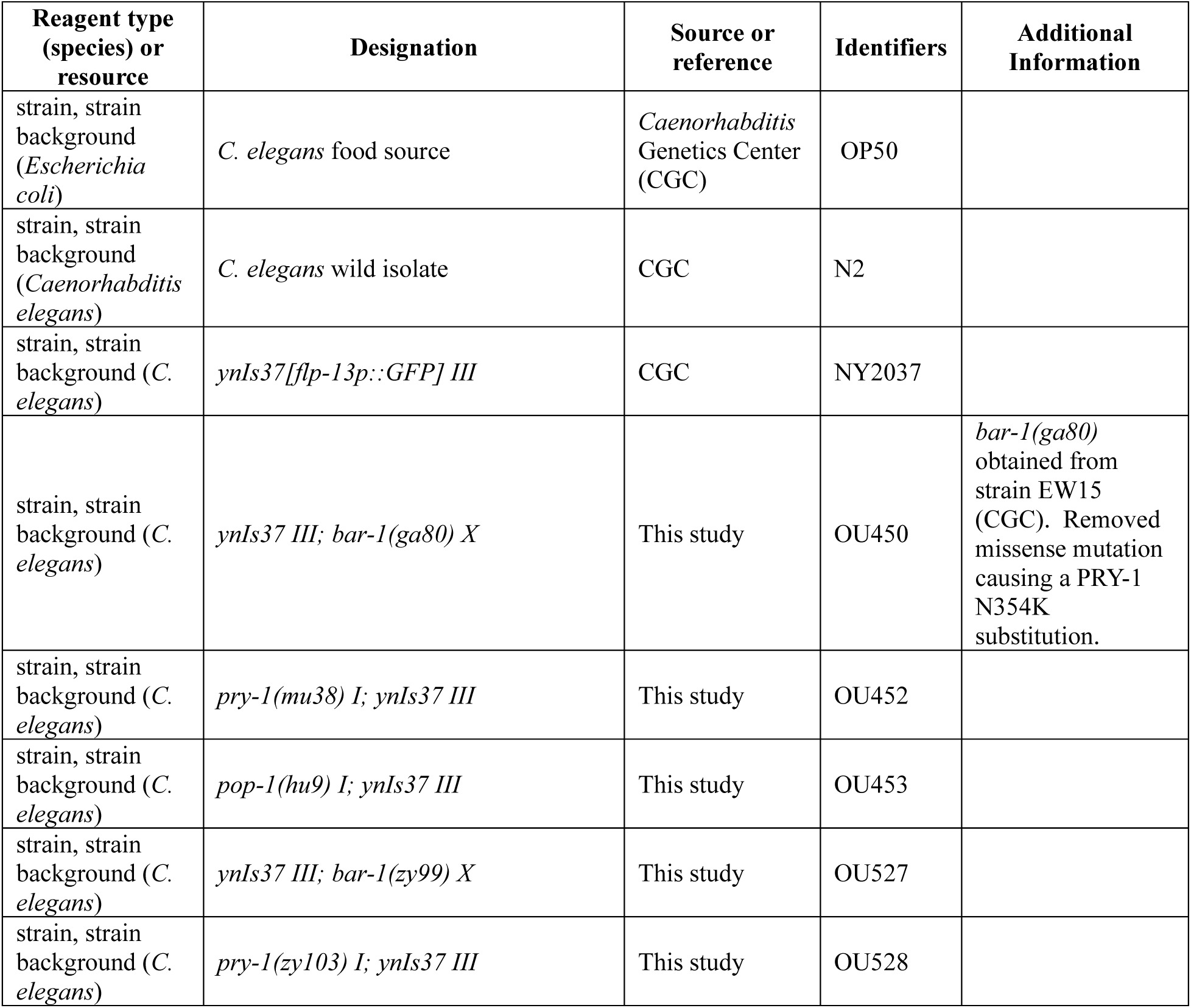
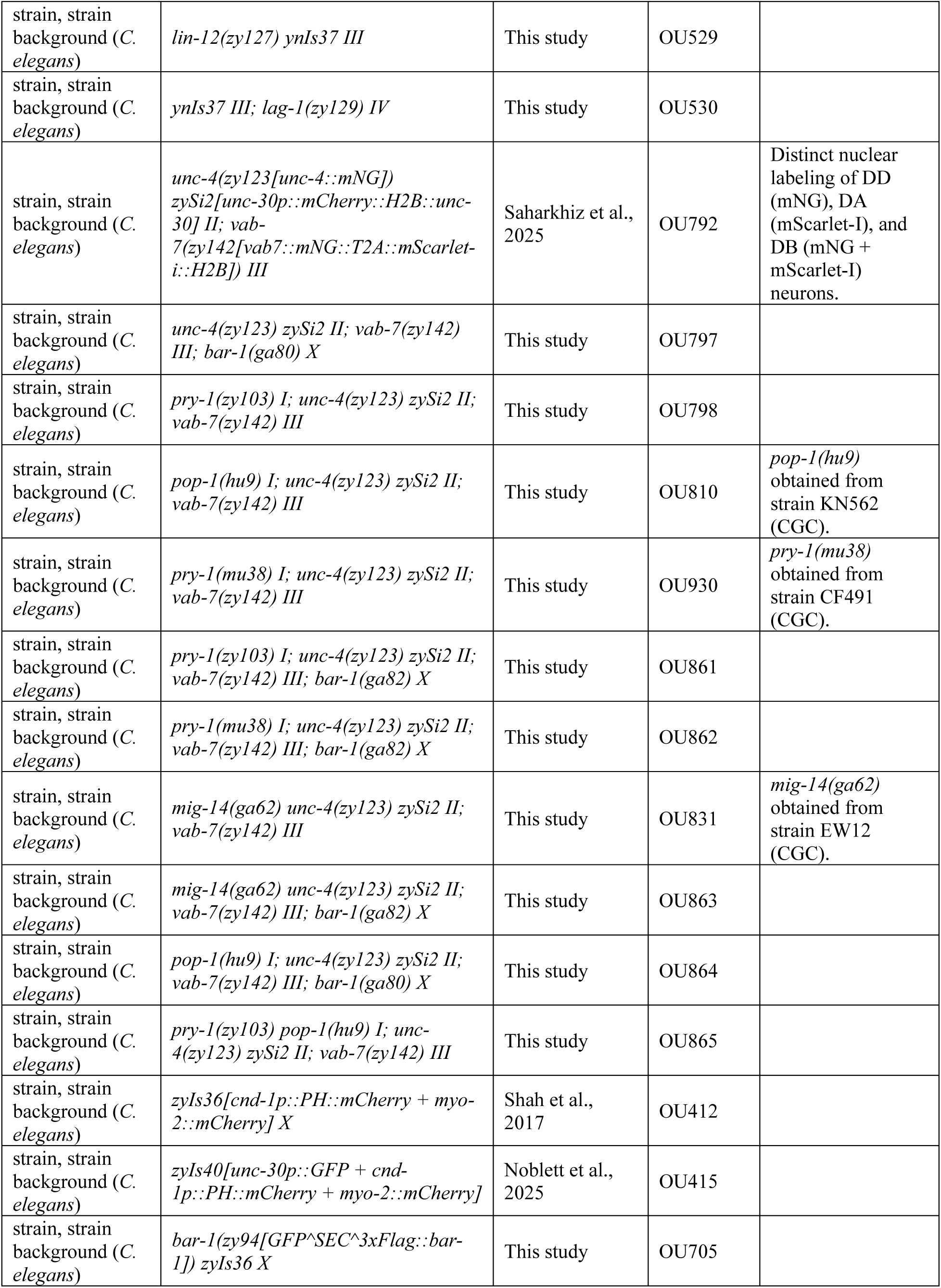
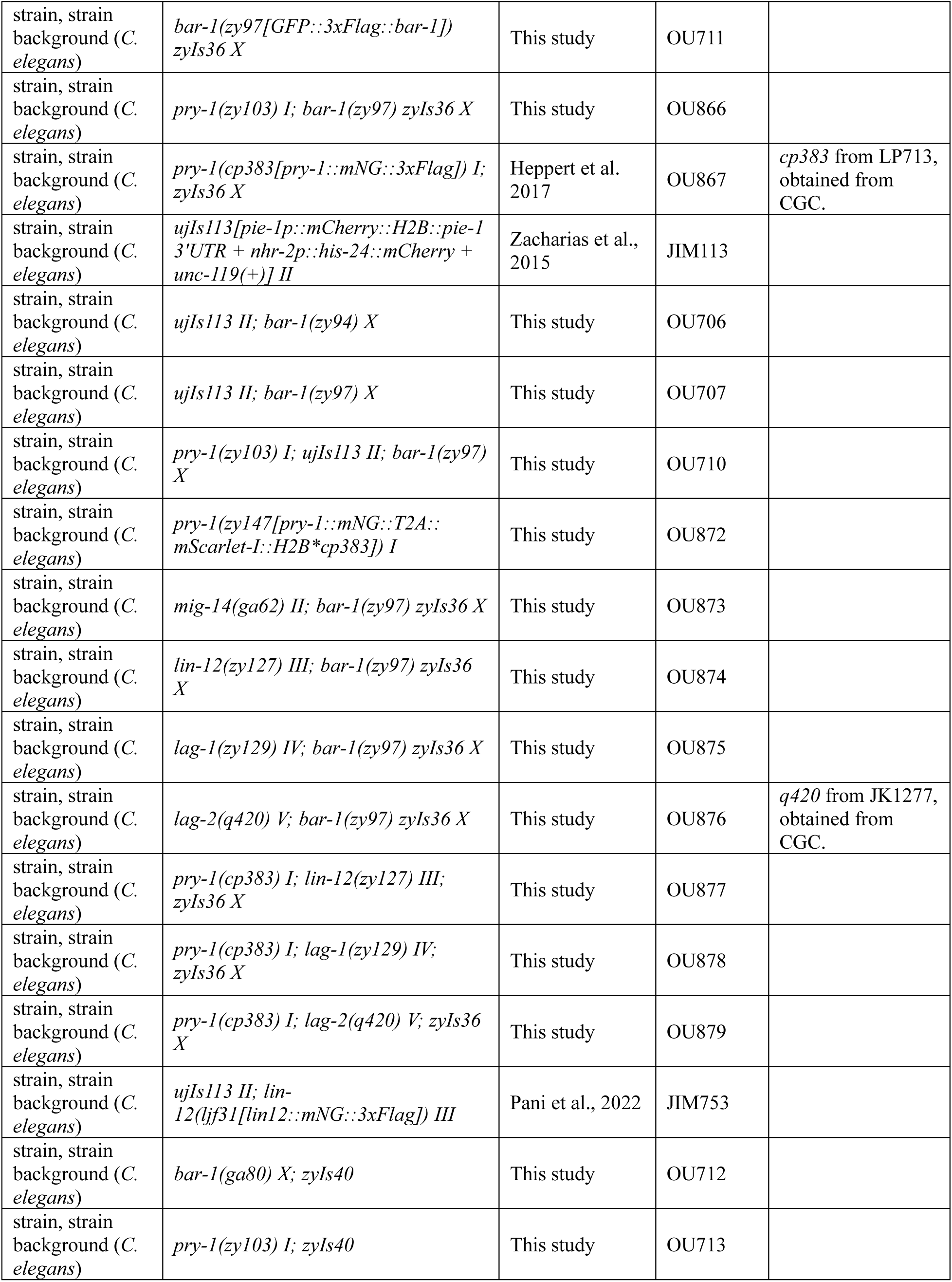
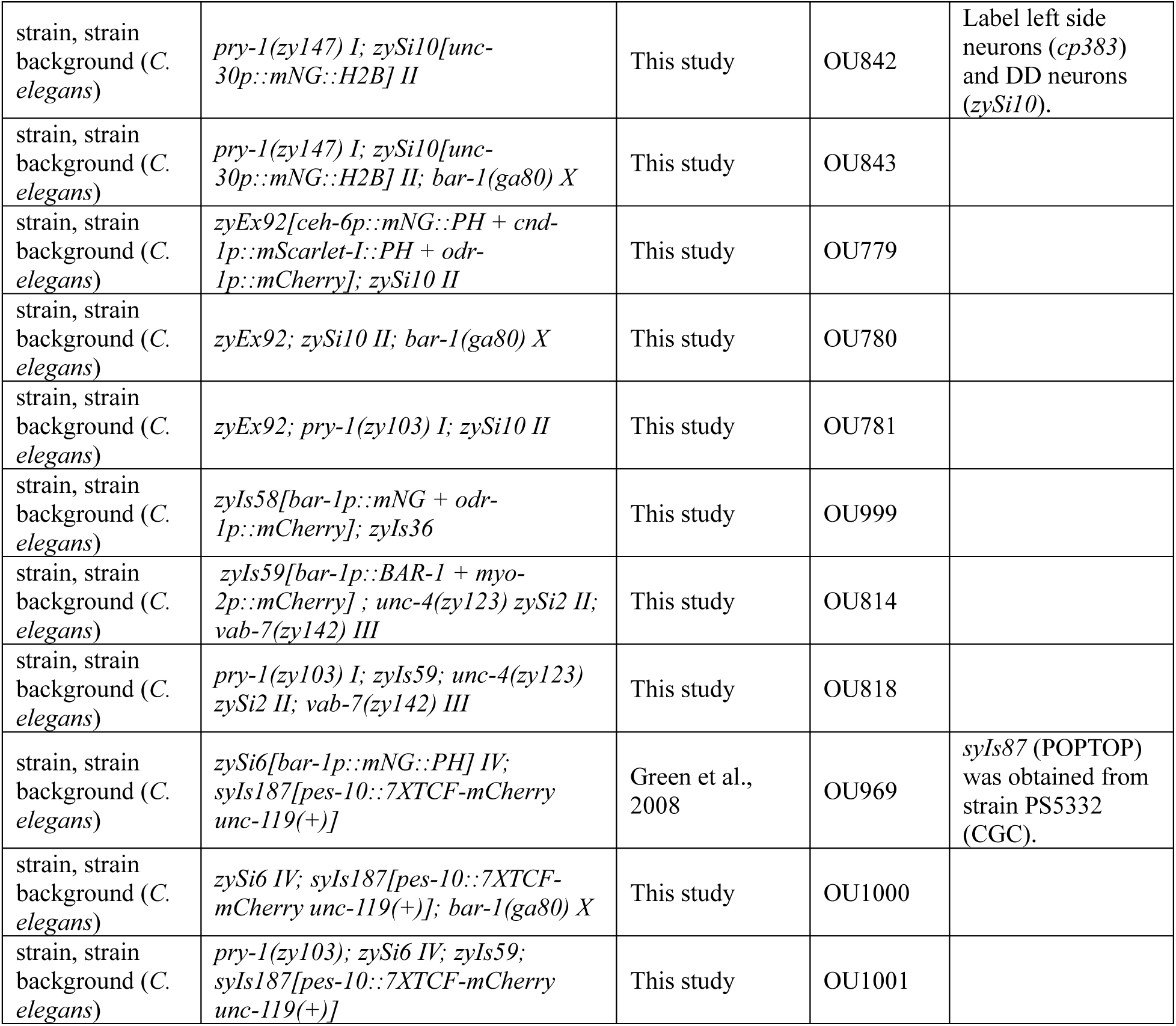
Key Resources Table.

## MATERIALS & METHODS

### *C. elegans* Strains and Alleles

*C. elegans* strains were cultured on standard NGM plates seeded with OP50 and maintained at 20°C. The Bristol N2 worm strain was used as the wild-type genetic background. Details for all strains and alleles used in this study as well as their sources are listed in the Key Resources Table.

### Motor neuron position measurements

The *unc-4(zy123[unc-4::mNG])*, *zySi2[unc-30p::mCherry::H2B]*, and *vab-7(zy142[vab-7::mNG::T2A::mScarlet-I::H2B])* reporters were used to label DA/SAB, DD, and DB motor neurons, respectively, in newly hatched L1 larvae as previously described by Saharkhiz et al. (2025). L1 larvae were mounted on 2% agarose pads and anaesthetized in a solution of 50 mM sodium azide in M9. Images used to analyze neuron position were acquired with a 40X/1.4 NA oil immersion objective and an Axiocam 702 mono camera mounted on a widefield Axio Imager.M2 widefield microscope (Zeiss).

DA, DB, and DD motor neuron positions along the VNC was scored relative to the anterior SAB neuron (0%) and rectum (100%) using the VNC-Dist software tool (Saharkhiz et al., 2025). Because individual neurons within the DA, DB, and DD classes could not be distinguished by our reporters in mutants with disrupted positioning, neurons were numbered according to their anterior-to-posterior position. For example, the most anterior DB neuron was designated DB1, followed by DB2, DB3 and so on, irrespective of their canonical wild-type identities. Custom R scripts using the ggplot2 package (Wickham, 2016) were used to generate violin and scatter plots showing neuron position and distribution. To assess cell order along the anterior-posterior axis, all 22 DA, DB, and DD neurons were ranked within each worm according to their VNC-Dist positions, from anterior to posterior, to generate a sequential order of neurons. The resulting ranked sequences were represented using custom graphs generated in R. Relative positions of each neuron, along with standard deviation, displacement from WT, Cohen’s d, effect size, and 95% confidence intervals of the mean for individual neuron subtypes in each genotype are summarized in Table S1.

Positions of DD2-DD6 in 1.5-fold and 2-fold stage embryos were measured relative to DD1 (0%) and the tail (100%). Measurements were acquired in Fiji by manually tracing along the VNC and tail using the Segmented Line tool (Schindelin et al., 2012). DD neurons and the developing VNC were labeled using *zyIs40[unc-30p::GFP + cnd-1p::PH::mCherry]*. Embryos were imaged as described above using a 63X Plan-Apochromat/1.4 NA oil immersion objective.

### DD spacing phenotype and VNC innervation gap defect correlation

The reporter transgene *ynIs37[flp-13p::GFP]* was used to label DD neuron cell bodies and projections to quantify innervation and positioning defects. Images of worms were acquired using the 40X oil-immersion objective on our Imager.M2 widefield setup. The Segmented Line tool in Fiji was used to measure the width of innervation gaps and distances between DD neurons along the VNC. A Spacing Index was used to quantify the extent of DD neuron spacing defect between DD1, DD2 and DD3. This index was calculated by dividing the distance from the cell bodies of DD1 to DD2 along the VNC by the distance from the cell bodies of DD2 to DD3. An index approaching 1 indicates relatively even spacing between DD1, DD2 and DD3, while an index approaching 0 indicates a shorter distance between DD1 and DD2 relative to the distance between DD2 and DD3. Innervation gap width was measured along the region of the VNC located between DD2 and DD3. Within this region, gap width was measured as the length of the segment that was absent of neuronal processes. To account for variation in innervation gap size caused by differences in worm sizing, we divided the raw gap width by the distance from the nerve ring to the rectum (as a proxy for body length) to obtain a relative measurement for innervation gap width.

### CRISPR-Cas9-mediated knock-ins in *bar-1* and *pry-1*

GFP was inserted at the N terminus of the endogenous bar-1 gene using the CRISPR/Cas9 self-excising drug selection cassette (SEC) approach described in Dickinson et al. (2015). An sgRNA targeting *bar-1* was cloned into the Cas9-sgRNA plasmid pDD162 (Addgene #47549) to generate pAC444. The homology directed repair plasmid (pAC445) was constructed using the GFP-SEC plasmid pDD282 (Addgene #66823). The resulting knock-in alleles were named *zy94[GFP::SEC::bar-1]* and, after SEC excision, *zy97[GFP::bar-1]*.

A *dpy-10* co-CRISPR approach (Arribere et al., 2014) was used to insert a T2A::mScarlet-I::H2B cassette between the mNG and 3’UTR in *pry-1(cp383[pry-1::mNG]* to generate *pry-1(zy147[pry-1::mNG::T2A::mScarlet-I::H2B*cp383])*. A PCR-based approach was used to fuse fragments amplified from genomic DNA containing the mNG::T2A::mScarlet-I::H2B cassette from *vab-7(zy142[vab7::mNG::T2A::mScarlet-I::H2B])* worms (Saharkhiz et al., 2025) and the *pry-1* 3’UTR from *cp383* worms (Heppert et al., 2018). The resulting PCR product was used as the DNA template for homology-directed repair. An sgRNA targeting a site within mNG was generated using pDD162 to generate pAC849. Oligonucleotide sequences are provided in Table S2.

### Generation of reporter and ectopic expression constructs and transgenic strains

The *bar-1p::mNG* transcriptional reporter (pAC495), containing an ∼1.9 kb internal deletion within the *bar-1* promoter, was generated using Gibson Assembly (Gibson et al., 2009). This reporter combines ∼3.7 kb of distal and 1.2 kb of proximal *bar-1* promoter sequences amplified from genomic DNA with the mNG coding sequence amplified from pDD268 (Addgene #132523), using pPD95.77 (a gift from A. Fire, Stanford) as the vector backbone. Germline transformation was performed using standard methods (Mello et al., 1991) by injecting 2.5 ng/μl of pAC495, 30 ng/μl of *odr-1p::mCherry*, and pBluescript (Stratagene) to a total DNA concentration of 100 ng/μl. The resulting extrachromosomal array was spontaneously integrated to generate the *zyIs58* line.

The *bar-1p::bar-1* ectopic expression construct (pAC544) was generated using Gibson Assembly by combining PCR-amplified fragments containing the *bar-1* promoter and vector backbone from pAC495, a *bar-1* cDNA from pDE248 (a gift from D. Eisenmann, University of Maryland), and the *tbb-2 3’UTR*. pAC544 was injected at 2.5 ng/μl, together with 2.5 ng/μl of *myo-2p::mCherry* (pCFJ90, Addgene #19327) and pBluescript to a total DNA concentration of 100 ng/μl. The resulting extrachromosomal array was integrated using UV irradiation (UVP CL-1000 cross-linker, 254 nm, 3.25×10^4^ µJ cm^-2^) to generate the *zyIs59* line.

The *bar-1p::mNG::PH* (pAC570) construct was generated using Gibson assembly by replacing the *mNG::unc-54 3’ UTR* cassette in pAC495 with an *mNG::PH::tbb-2* 3’UTR cassette. A single copy of the *bar-1p::mNG::PH* reporter was then inserted near the *cxTi10882* MosI landing site on LG IV (Frøkjær-Jensen et al., 2008) using a SEC-mediated CRISPR/Cas9 approach to generate *zySi6[bar-1p::mNG::PH]*. Briefly, Gibson assembly was used to generate a homology-directed repair plasmid containing *bar-1p::mNG::PH* (derived from pAC570), a SEC cassette flanked by loxN sites (derived from pJH126; Addgene #172147) and approximately 700 bp homology arms flanking the insertion site. An sgRNA targeting a site adjacent to *cxTi10882* was generated as described above (pAC685), using the oligonucleotide listed in Table S2.

*cnd-1p::mScarlet-I::PH* (pAC740) was generated by Gibson assembly of an approximately 3.2 kb *cnd-1* promoter fragment, an mScarlet-I fragment from pGLOW39 (Addgene #173068), and a PH::tbb-2 3′ UTR-pPD95.77 fragment. *ceh-6p::mNG::PH* (pAC795) was generated as described for pAC495, except the bar-1 promoter was replaced with an approximately 6.2 kb of ceh-6 promoter fragment amplified from genomic DNA. The *ceh-6* promoter was used to label DB neurons based on previous expression studies showing *ceh-6* expression in DB1-7, in addition to their sister cells and a relatively small number of other cells (Sarov et al., 2012). *zyEx92* was generated by injecting 0.5 ng/μl of pAC740, 1.5 ng/μl of pAC795, 30 ng/μl of *odr-1p::mCherry*, and pBluescript to a total DNA concentration of 100 ng/μl.

The *unc-30p::mNG::H2B* (pAC776) construct was generated using Gibson Assembly by combining PCR-amplified fragments containing the *unc-30* promoter, *mNG*, *H2B::tbb-2 3’UTR*, and the vector backbone from constructs described in Saharkhiz et al. (2025). *zySi10[unc-30p::mNG::H2B]* was generated using the same approach as *zySi6*, but using *ttTi5605* on LGII as the landing site, approximately 700 bp homology arms flanking the site, and an sgRNA targeting a site adjacent to *ttTi5605*. The sgRNA (pAC670) was generated using the oligonucleotide listed in Table S1.

### Confocal microscopy

Imaging of *zy94[GFP::SEC::bar-1]*, *zy97[GFP::bar-1]*, *pry-1(cp383[pry-1::mNG]*, and *pry-1(zy147[pry-1::mNG::T2A::mScarlet-I::H2B*cp383])* for localization analysis, and of POPTOP for fluorescence intensity quantification, was performed using an EM-CCD camera (Hamamatsu ImagEM) mounted on a Leica DMI6000B inverted microscope stand with an attached Quorum spinning disk. Embryos were mounted on a 2% agarose pad in a 50 mM solution of sodium azide to immobilize twitching embryos. Fluorophores were excited using an X-Cite 120 Mercury light source. Images were acquired using a 63X/1.4 NA oil immersion objective. For representative images of the *bar-1* and *pry-1* fluorescent reporters, excitation intensity and exposure time were adjusted differently for each reporter because expression levels varied considerably.

Standardized imaging settings were used to quantify POPTOP fluorescence intensity. POPTOP was excited using a 561nm laser at 75% intensity with an exposure time of 223 ms. The VNC marker *zySi6[bar-1p::mNG::PH]* was excited using a 490 nm laser at 75% intensity with an exposure time of 180 ms. Z-stacks were acquired at 0.5 µm intervals, covering the volume of the VNC, with the start and stop z-positions adjusted for each embryo.

### Lineage tracing

Embryos were mounted for imaging as in (Bao and Murray, 2011) and imaged at 22°C using a Leica TCS SP5 or Stellaris resonance scanning confocal microscope (67 z planes at 0.5 μm z-spacing and 1.5 minute time spacing), as described in (Murray et al., 2022). We used StarryNite (Santella et al., 2010) to trace lineages and the AceTree suite (Boyle et al., 2006) to correct errors in the automated analysis and quantify gene expression. For *zy94[GFP::SEC::bar-1]* and *zy97[GFP::bar-1]* expression we used the ‘blot’ (Murray et al., 2008) approach to correct for local background in nuclear protein quantification by subtracting nearby non-nuclear signal. For LIN-12::mNG, we used raw intensity (‘none’ correction method from Murray *et al* 2008) because the remaining membrane-localized LIN-12 signal otherwise masks changes in nuclear LIN-12. However, both reporters give qualitatively similar results to the other background correction approaches.

### Lightsheet microscopy and analysis

Lightsheet microscopy was performed using an ASI diSPIM equipped with symmetric Nikon CFI Apo 40XW NIR objectives. Illumination was provided by an Oxxius L4Cc laser combiner with 488 and 561 nm laser modules. Dual color imaging was performed simultaneously using a Hamamatsu W-View Gemini image splitter and acquired using a Hamamatsu Orca Fusion C14440-20UP camera. Embryos were collected by microcapillary pipette from NGM plates, washed with M9 to remove excess bacteria and mounted in a diSPIM sample holder onto a 25×50 mm coverslip which had previously been coated with poly-l-lysine and allowed to dry. Embryos were imaged from a single view using 1 μm steps in Z using Micro-Manager 1.4 and the ASI diSPIM plugin to control acquisition at a time interval of 2 minutes between t-slices.

The membranes of the developing DA (excluding DA6, DA7, and DA9), DB and DD motor neurons were labeled using *zyEx92[ceh-6p::mNG::PH + cnd-1p::mCherry::PH]* and the nuclei of DD neurons labeled with *zySi10[unc-30p::mNG::H2B].* Each timelapse was aligned in the time dimension by observing the cell divisions which generate DD2/DD4, DD3/DD5, DA2/DA4, and DA3/DA5 (marked by *cnd-1p::mCherry::PH*) and AIAL/DB6 and AIAR/DB7 (marked by *ceh-6p::mNG::PH*). The completion of the division that generates the last of these cells is labeled as 0 mins. Timelapses were followed forward at each t-slice to observe the state of DD1-DD2 separation. The first time point at which the anteroposterior-oriented DD1-DD2 was clearly separated by an intervening junction formed between two or more medial-lateral cells (excluding when DD1 and DD2 meet at a vertex) was considered the time point at which DD1-DD2 was considered as ‘separated’; it was verified that after this timepoint, DD1 and DD2 never reforms a junction at any point. If DD1-DD2 did not separate by the end of the timelapse, the state of DD1-DD2 was categorized as ‘unseparated’. Beginning at 1.5-fold stage, neurons were identified at each timeframe by assigning cells in decreasing numerical order, from posterior to anterior of the VNC, according to the expected WT order of neurons. Neurons expressing only *cnd-1p::mCherry::PH* were labelled DA5 to DA1. Neurons labelled with both *cnd-1p::mCherry::PH* and the nuclear *zySi10* mNG reporter were labelled DD6 to DD1. Neurons that were only marked by *ceh-6p::mNG::PH* were labelled DA8, DB7 to DB4, and then DB2. Lastly, neurons labelled by both *ceh-6p::mNG::PH* and *cnd-1p::mCherry::PH* were labelled DB3 followed by DB1. To highlight the progression of neuron movement in each strain, select timepoints were traced from a representative timelapse to show neuron shape and position in each fluorescence image; due to embryonic twitching, the orientation of the VNC varied across timepoints in these tracings. To simplify the visualization and facilitate readability, tracings were simplified and rotated (if needed) to generate illustrations of cell positions oriented in the same ventral viewpoint.

### POPTOP fluorescence intensity measurement

Confocal spinning disk images were used to quantify POPTOP fluorescence intensity in embryos. Intensity measurements were performed using a custom analysis pipeline based in Fiji/ImageJ 2.14.0/1.54f software (Schindelin et al., 2012). The fluorescence signal from *zySi6[bar-1p::mNG::PH]* was used to segment the developing VNC. For measurements at 2-fold stage, a segmentation classifier was generated using the Trainable Weka Segmentation plugin (v3.3.4) (Arganda-Carreras et al., 2017) and trained on the *zySi6* signal. The classifier was then used on z-stack images for each sample to output a segmented image which was then used to generate unique regions-of-interest (ROI) covering the area corresponding to the VNC for each z-slice. Segmentation quality was visually inspected for accuracy before proceeding with measurements. We then used these unique ROIs at each z-slice in the POPTOP channel to measure ‘Area’ and ‘Integrated Density’. A raw mean fluorescence intensity (MFI) measurement for any given sample was calculated by dividing the summation of integrated densities (IntDen) by the summation of area as measured in each z-slice ROIs. To estimate the amount of signal corresponding to background noise, we manually drew an area outside the embryo and performed the same MFI measurement per z-slice per sample. The final MFI measurement used for statistics and plotting was calculated by subtracting background MFI from the raw MFI.

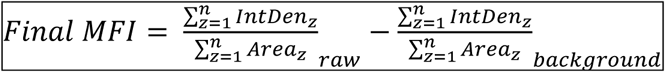

Where *n* is the total number of z-slices in the analyzed stack, *z* is the *z-slice* index.

POPTOP measurements for bean stage embryos were performed using a modified procedure since Weka segmentation did not allow for the separation of the left from the right VNC. For each z-stack image, a range of z-slices covering the volume of the VNC was selected and used to produce a Sum Slices z-projection. The Segmented Line tool was then used to manually segment the area corresponding to the left or right VNC (using the *zySi6* fluorescence channel as reference) to obtain an area measurement and a POPTOP integrated density measurement. Background noise measurement and final MFI calculation were performed in the same way as described above keeping the left and right measurements separate. To obtain the POPTOP fluorescence for the total VNC area, we added the left and right measurements from Fiji prior to calculating the final MFI.

### Statistics

Statistical analyses were performed using R version 4.3.3 and RStudio. Graphs were generated using the ggplot2 package (Wickham, 2016).

Relative positions of DA, DB, and DD neurons at L1 were extracted from VNC-Dist. Mean relative positions of individual neurons in each mutant were compared with the WT control using Welch’s t-tests. P-values from these comparisons were adjusted for multiple comparisons using Benjamini-Hochberg’s false discovery rate (FDR) correction. Adjusted p-values were used to annotate the mean relative position graphs (also provided in Table S1). To assess genetic interactions, relative positions of individual neurons were compared among selected mutant strains, including WT, by using an ANOVA test. When appropriate, Tukey’s honest significant difference (HSD) post-hoc test was performed to compare mean neuron positions between pairs of strains. Comparisons performed this way are presented as violin plots.

The Spacing Index and normalized innervation gap widths were compared between WT and *bar-1(ga80)* larvae using Welch’s t-test. To assess the correlation between innervation gap width and the Spacing Index, we pooled the samples from WT and *bar-1(ga80)* mutants (which provided better coverage of the range of Spacing Index scores) and performed Pearson’s correlation analysis.

To compare DD positioning in 1.5-fold and 2-fold embryos expressing *zyIs40*, we performed a one-way ANOVA with Dunnett’s post-hoc test to compare the positioning of each DD neuron among WT, *bar-1(ga80)* and *pry-1(zy103)* mutants.

POPTOP expression intensities were compared using Welch’s t-test to compare the final MFI. Comparisons included POPTOP levels between the VNC cells of WT and *zyIs59; pry-1(zy103)* 2-fold embryos, between left and right VNC cells of WT bean stage embryos, between left and right VNC cells of *bar-1(ga80)* bean stage embryos, and between VNC cells of WT and *bar-1(ga80)* bean stage embryos.

To compare the frequency of DD1-DD2 order phenotypes between WT or *bar-1(ga80)* 2-fold embryos labeled with *zy147; zySi10*, we performed Fisher’s exact test on a 2 genotypes x 2 phenotypes contingency table. The ‘Anterior’ group includes embryos where DD1 is anterior to DD2; the ‘Posterior’ group includes embryos where DD1 is posterior to DD2. DA2-DD2 neuron order was compared using Fisher’s exact test on a 2 genotypes x 3 phenotypes contingency table. The ‘Anterior’ and ‘Posterior’ phenotypes are as defined above with DA2 instead of DD1. Embryos where the nuclei of DA2 and DD2 overlapped (when viewed from the left or right side of the embryo) were categorized as a third phenotypic group labelled ‘Unintercalated’.

## RESULTS

### β-catenin-TCF signaling, negatively regulated by Axin, is required for proper motor neuron cell body positioning

The stereotyped pattern of DD, DA and DB motor neurons is assembled beginning at the embryonic bean stage, when post-mitotic neurons born on the left and right sides migrate toward the midline and intercalate to form an ordered linear arrangement by the 2-fold stage (Shah et al., 2017). We previously showed that *vang-1*/VANGL and *prkl-1*/PRICKLE, components of the non-canonical Wnt planar cell polarity pathway, are required during these initial stages to assemble these neurons into their proper positions within the VNC (Shah et al., 2017). These findings led us to investigate whether β-catenin-TCF-dependent Wnt signaling also contributes to motor neuron positioning. *bar-1* is one of four β-catenin genes in *C. elegans* and acts as a transcriptional co-activator with *pop-1*/TCF (Korswagen et al., 2000; Robertson and Lin, 2012), whereas *pry-1*/Axin negatively regulates BAR-1-dependent Wnt signaling (Gleason et al., 2002; Korswagen et al., 2002). To assess their roles, we used the *ynIs37[flp-13p::GFP]* transgene to label DD neurons in the *bar-1(ga80)* and *pry-1(mu38)* null mutants and the *pop-1(hu9)* hypomorphic mutant. All three mutants exhibited DD positioning defects, primarily affecting DD1 and DD2 (Fig. S1). In *bar-1* and *pop-1* mutants, DD2 shifted anteriorly, whereas in *pry-1* mutants, DD1 shifted posteriorly compared to their positions in WT. These phenotypes contrasted with the effects of *vang-1* and *prkl-1* loss, which caused nearly all DD neurons to shift anteriorly (Shah et al., 2017).

To quantify DD position shifts and to determine whether DA and DB neurons were also affected in *bar-1*, *pop-1*, and *pry-1* mutants, we used VNC-Dist, a Python-based software tool we developed to map relative positions of motor neurons along the VNC from annotated images of L1 stage larvae (Saharkhiz et al., 2025). This tool utilizes a two-color fluorophore scheme to distinctly label motor neuron nuclei with class-specific markers: DD1-6 (a single copy insertion of *unc-30p::mCherry::H2B*), DA1-9 along with SAB neurons in the retrovesicular ganglion (endogenously tagged *unc-4::mNG*), and DB1-7 (endogenously tagged *vab-7::mNG::T2A::mScarlet-I::H2B*) (Fig. 1A). VNC-Dist measures the position of each motor neuron nucleus along the ventral curvature of the worm body relative to two reference points to accommodate for variations in worm length: the most anterior SAB neuron (0%) and the rectum (100%). Using the relative position output for each neuron in each strain as our data points, we performed several analyses to compare strains. We first compared our mutant strains against the WT control to determine whether mean neuron positions are shifted from the WT position (Fig 1B and Table S1). Next, we generated violin plots to visualize the distribution of select neurons and compare between mutant strains to test for genetic interactions and epistatic relationships (Fig. 1C, D and E). Lastly, we used the positional information of each neuron within each worm to generate plots that display the relative order of DA, DB, and DD neurons, enabling visualization of patterning in key mutants compared to WT (Fig. 1F). For these analyses, we also included *pry-1(zy103)*, a new allele isolated from a genetic screen for DD neuron positioning defects (unpublished) that carries a premature stop codon (R355stop; Table S3). Unlike *pry-1(mu38)* (W268stop), *pry-1(zy103)* does not show a severe reduction in viability. In subsequent analyses, we used only the *pry-1(zy103)* mutant, as it produces motor neuron positioning defects similar to those observed in *pry-1(mu38)* but has much higher viability.

**Figure 1.**
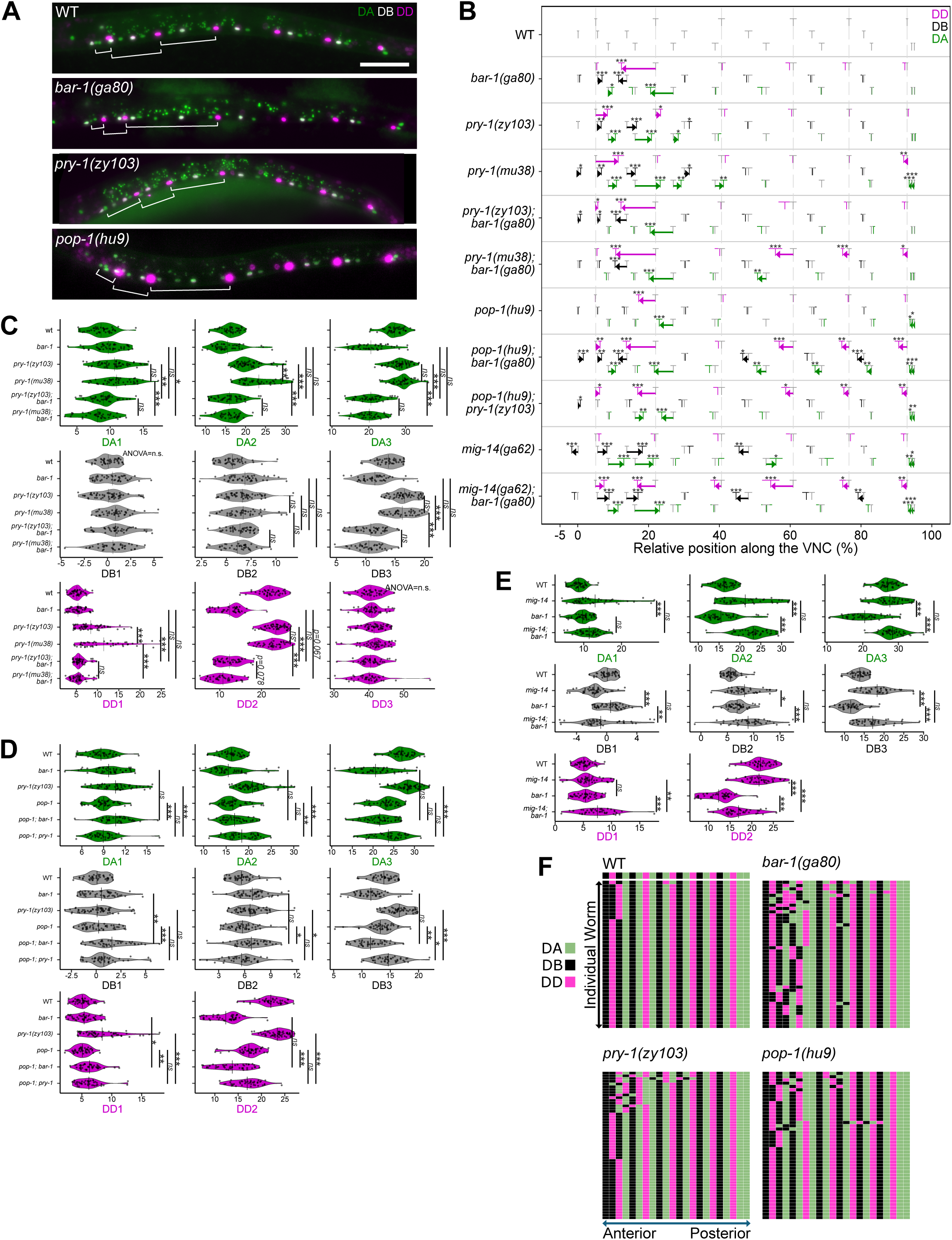
BAR-1/β-catenin and PRY-1/Axin regulate motor neuron cell body position in the VNC. (A) Representative fluorescence images of L1 worms showing the positions of DD, DA, and DB motor neuron cell bodies in WT and *bar-1*, *pry-1*, and *pop-1* mutants. DD, DA (and SAB), and DB neuron nuclei are labeled with mCherry (magenta), mNG (green), and mNG/mScarlet (magenta + green, pseudocolored white), respectively. Brackets indicate the spacing between the first SAB to DD1, DD1 to DD2, and DD2 to DD3 neuron cell bodies to highlight positional shifts between genotypes. Scale bar, 100 µm. (B) Mean relative position and 95% confidence interval of neurons in WT and various mutant backgrounds. Position is expressed as a percentage of total VNC length, measured from the most anterior SAB neuron (0%) to the rectum (100%). Arrows indicate the direction and magnitude of statistically significant shifts in mean position from WT to mutant neurons with moderate or greater effect sizes (|Cohen’s d| > 0.5). Dotted vertical lines indicate the mean position of WT DD neurons for visual reference. Sample sizes as well as other statistical values are listed in Table S1. Mutant positions were compared to WT using Welch’s *t*-test followed by multiple comparisons correction using the Benjamini-Hochberg procedure for controlling the false discovery rate to obtain adjusted *p* values (*p<0.05, **p<0.01, ***p<0.001). (Table S1). (C-E) Violin plots showing the distribution of individual cell body position data from (B) for a subset of anterior DD, DA, and DB neurons. Each data point represents a single neuron from a single animal. To test genetic interactions, we identified differences in position among selected mutants and WT by performing an ANOVA followed with Tukey’s HSD post-hoc test. The significance levels of only the biologically relevant comparisons are shown. Each figure panel represents a different group of mutants analysed statistically, with the exception of (D) which shows statistics comparing the *pop-1; bar-1* double mutant with the corresponding single mutants, as well as a separate statistical analysis comparing the *pop-1; pry-1* double mutant with the corresponding single mutants. (F) Maps showing VNC motor neuron order in WT and *bar-1*, *pry-1*, and *pop-1* single mutants, generated from the positional data in (B). Each map shows the relative anteroposterior arrangement of DD (magenta), DA (green), and DB (black) neurons along the VNC. The most common WT neuronal order is shown as a separate strip above the WT map for reference.

As observed in DD neurons, we found that the β-catenin pathway mutations shift the mean AP position of a subset of DA, DB, and DD neurons that are primarily located in the anterior VNC. *bar-1* and *pop-1* mutants primarily exhibit anterior-displaced neurons, with DD2 and DA3 showing the most pronounced shifts (all p < 0.001, Fig. 1A, 1B and Table S1). In contrast, *pry-1* mutants displayed posterior-displaced neurons, particularly DD1, DA2, and DB3 (all p < 0.001), with *pry-1(mu38)* showing a generally stronger phenotype consistent with it being a null allele. These position defects were accompanied by changes in the relative order of motor neurons, particularly in the anterior region of the VNC (Fig. 1F).

We also examined double mutants to investigate genetic interactions between *bar-1(ga80)*, *pry-1(zy103)*, and *pop-1(hu9)*. As expected, given Axin’s role in the β-catenin destruction complex, mean shifts and individual neuron distributions in *pry-1; bar-1* double mutants resembled those of *bar-1* single mutants, and *pop-1; pry-1* double mutants resembled *pop-1* single mutants (Fig. 1C and D, Table S4 and S5). This indicates that *pry-1* acts upstream of both *bar-1* and *pop-1*. Specifically, the DA2 (p < 0.001), DB3 (p < 0.001), and DD1 (p < 0.001) position defects in *pry-1* mutants are suppressed in *pry-1; bar-1* double mutants. In contrast, the anteriorly displaced DA3 (p = 1.00) and DD2 (p = 1.00) defects in *bar-1* mutants are unaltered in *pry-1; bar-1* double mutants. The neurons that shifted most in the *bar-1* and *pop-1* single mutants showed similar mean shifts in *pop-1; bar-1* double mutants (Fig. 1D and Table S6). DA3 position in *pop-1; bar-1* was not significantly different compared to *bar-1* (p = 0.486) or *pop-1* (p = 0.175). DD2 position in *pop-1; bar-1* was not significantly different compared to *bar-1* (p = 0.119) and, although significantly more anteriorly shifted compared to *pop-1* (p < 0.001), occur in the same direction relative to WT. These phenotypes are consistent with BAR-1 and POP-1 acting together in the same pathway. Interestingly, we also observed additional shifts in posterior VNC neurons in *pop-1; bar-1* double mutants that were absent or much less pronounced in the single mutants, suggesting that *bar-1* and *pop-1* also act in posterior neurons in a partially redundant manner (Fig 1B). Together, these findings indicate that *pry-1* acts in its canonical role as a negative regulator of *bar-1,* and consequently the BAR-1/POP-1 complex, to regulate motor neuron positioning in the anterior VNC.

### Motor neuron position defects in *bar-1* mutants correlate with innervation gaps

The cell bodies of the 6 DD neurons are spaced relatively evenly along the VNC. Each DD neuron extends processes anteriorly along the ventral cord which then continue in the dorsal cord via commissures (White et al., 1976). This arrangement enables DD neuronal processes to extend along the length of the ventral and dorsal cords, providing innervation of body wall muscles required for sinusoidal movement. We therefore asked whether the anterior displacement of DD2 in *bar-1* mutants (Fig. 1), which brings DD2 closer to DD1 and increases the DD2-DD3 distance, affects innervation. *ynIs37[flp-13p::GFP]* was used to visualize the cell bodies and processes of DD neurons in L1 larvae. While most WT animals had little or no innervation gaps, *bar-1(ga80)* mutants displayed regions along the ventral and dorsal nerve cords that lacked DD processes (Fig. 2A). Quantification of the gap between DD2 and DD3 showed that *bar-1* mutant larvae (n = 15) exhibited significantly wider gaps compared to WT (WT n = 15, *bar-1* n = 15; p < 0.001) (Fig. 2B and 2C), indicating that the innervation of body wall muscle by DD2 is disrupted in *bar-1* mutants.

**Figure 2.**
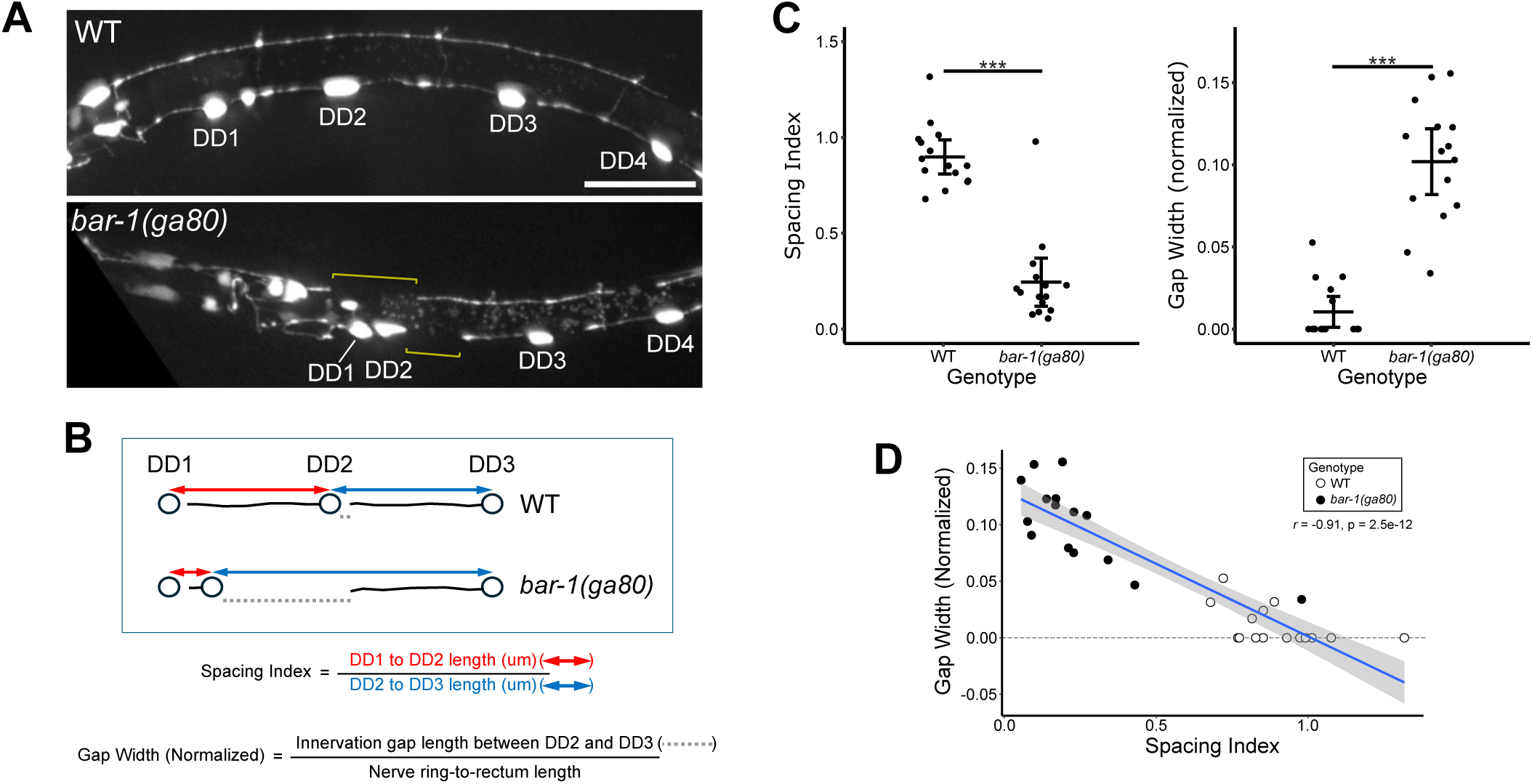
Motor neuron cell body displacement in *bar-1* mutants is associated with axon innervation gaps in L1 larvae. (A) Representative images of WT and *bar-1(ga80)* L1 larvae expressing *ynIs37[flp-13p::GFP]*, which labels DD neurons and their projections. Brackets highlight innervation gaps in the ventral and dorsal nerve cord of *bar-1* mutants. Scale bar, 100 µm. (B) Diagram showing the measurements used for the calculation of the spacing index (a quantification of the evenness of the spread in cell body positions of DD1, DD2, and DD3) and the normalized gap width (a measure of the length of the gap in innervation in the VNC region located between DD2 and DD3) (C) Dot plots comparing the spacing index and normalized gap width of WT and *bar-1* larvae. Measurements were compared using Welch’s t-test ***p < 0.001. Sample sizes were n = 15 for WT and n = 15 for *bar-1(ga80)*. (D) Scatter plot showing the correlation between the innervation gap width and spacing index among the pooled WT and *bar-1*datasets (n = 30 total). Each data point represents a single animal. Pearson’s correlation coefficient was used to assess the relationship between the two variables.

Because the DD2 cell body is significantly displaced anteriorly in *bar-1* mutants, we asked whether the altered spacing between DD1, DD2, and DD3 is associated with the width of the DD2-DD3 innervation gap. To quantify the extent of altered DD neuron spacing, we developed a Spacing Index calculated as the distance between the cell bodies of DD1 and DD2 divided by the distance between DD2 and DD3 (Fig. 2B). An index score approaching 1 indicates relatively even spacing, whereas an index below or above 1 indicates relative unevenness. *bar-1* mutant animals exhibited a significantly lower Spacing Index than WT animals (p < 0.001) (Fig. 2C). In WT larvae, the distance between DD1 and DD2 was on average 0.90 times the distance between DD2 and DD3. In *bar-1* mutants, it was 0.24 times the distance between DD2 and DD3. We next asked whether the extent of altered DD spacing was related to the severity of the innervation gap. Pearson’s correlation analysis of the pooled WT and *bar-1* datasets (combined n = 30) revealed a strong linear correlation between the DD2-DD3 innervation gap width and the Spacing Index (*r* = -0.91, *p < 0.001*) (Fig. 2D), showing that wider innervation gaps were associated with more uneven DD neuron spacing. These findings suggest that motor neuron cell body position is important for proper connectivity, and that AP shifts in cell body positions can contribute to local innervation defects.

### BAR-1 likely acts as a negative regulator of a MIG-14/Wntless-dependent pathway

Secreted Wnt proteins activate the canonical pathway by binding to Frizzled receptors, leading to the release of β-catenin from Axin-mediated repression (Nusse and Clevers, 2017). The *C. elegans* genome encodes five Wnt ligands (Korswagen, 2002), raising the possibility of functional redundancy among them. To perturb Wnt secretion broadly and thereby circumvent potential redundancy among individual Wnt ligands, we examined *mig-14(ga62)*, a hypomorphic allele of the *C. elegans* homolog of Wntless. MIG-14 is an essential membrane protein required for Wnt secretion and its disruption is expected to impair secretion of all Wnt ligands (Bänziger et al., 2006; Nusse and Clevers, 2017). We used this hypomorphic allele because complete loss of *mig-14* is lethal. Mutations in *mig-14* produce a range of phenotypes that resemble those caused by disruption of specific Wnt signaling pathways (Pan et al., 2008; Yang et al., 2008).

We therefore asked whether reduced *mig-14* function produces motor neuron positioning defects like those found in *bar-1* or *pry-1* mutants. *mig-14* mutants predominantly displayed posterior displacements, in contrast to the anterior shifts observed in *bar-1* mutants (Fig 1B). Additionally, unlike *bar-1* mutants, DD2 position was unaffected in *mig-14* mutants. Position shifts in *mig-14* mutants occurred primarily in DA and DB neurons (most notably DA1, DA2, DB2, and DB3; all p < 0.001) (Fig. 1B and Table S1). Because Wnt signaling typically positively regulates BAR-1 activity, we expected that *mig-14* mutants would phenocopy *bar-1* loss of function, however, *mig-14* mutants displayed a largely distinct phenotype affecting different neurons. To further characterize the relationship between BAR-1 and Wnt signaling, we analyzed neuron position defects in *mig-14; bar-1* double mutants (Fig. 1B and 1E, Table S7). We found that the DA and DB neuron defects observed in *mig-14* single mutants (DA1 (p = 0.184), DA2 (p = 0.105), DB1 (p = 0.138), DB2 (p = 0.558), and DB3 (p = 0.349)) persisted in the double mutants. In contrast, the DD2 (p < 0.001) and DA3 (p < 0.001) positional shifts observed in *bar-1(ga80)* single mutants were, respectively, partially and fully suppressed. These results suggest that *mig-14* is epistatic to *bar-1* during VNC neuron positioning, consistent with *mig-14* or a *mig-14*-dependent pathway acting downstream of *bar-1*. Interestingly, *mig-14; bar-1* double mutants displayed position defects in a subset of additional neurons not observed in either single mutant, most notably DD4 (p < 0.001 compared to WT), suggesting that *bar-1* and *mig-14* may act redundantly to regulate the position of these neurons. Thus, while Wnt ligands contribute to neuron positioning, particularly DA and DB neurons, our results are inconsistent with the canonical model in which Wnt positively regulates BAR-1 to establish motor neuron position.

### BAR-1 protein is asymmetrically expressed in right-side neurons during VNC assembly

To better understand how *bar-1* controls motor neuron positioning, we examined the timing and spatial pattern of *bar-1* expression during VNC assembly. To visualize BAR-1 expression at single cell resolution, we used a self-excising cassette (SEC)-mediated CRISPR/Cas9 approach to insert GFP at the N-terminus of the endogenous *bar-1* gene (Fig. 3A) (Dickinson et al., 2015). This method resulted in two lines, *zy94[GFP::SEC::bar-1]* and, after SEC excision, *zy97[GFP::bar-1]*. Because the SEC cassette in *zy94* introduces a stop codon upstream of the *bar-1* start codon, GFP fluorescence indicates *bar-1* promoter activity rather than protein expression, allowing us to distinguish endogenous transcription (*zy94*) from protein (*zy97*) expression. Expression patterns were examined in combination with *zyIs36[cnd-1p::PH::mCherry]*, a transgene that labels neuronal membranes, including those of DD1-6, DA1-5, DB1, and DB3 (Fig. 3B-D). In *zy94*, GFP fluorescence driven by the *bar-1* promoter was first detected at the bean stage in DD and DA neurons co-expressing membrane-localized mCherry after the terminal divisions that give rise to individual motor neurons (n > 30). Interestingly, *bar-1* activity was initially observed in the more posterior DD3-6 and DA2-5 neurons (Fig. 3B), before expanding anteriorly to include DD1-2 and DA1 by late bean/early comma stage. GFP expression in these neurons persisted into the 1.5-fold stage (Fig. 3D) and continued through to the larval stage (n > 30). Strikingly, when we examined endogenous GFP::BAR-1 protein expression in *zy97* (n > 30), GFP fluorescence was found exclusively in right-side derived DD and DA neurons, where it was predominantly localized to the nucleus, following the same posterior to anterior temporal progression as *bar-1* promoter activity (Fig. 3B and 3C). These observations indicate that endogenous BAR-1 is asymmetrically restricted to right-side neurons during VNC assembly, despite transcriptional activity on both sides.

**Figure 3.**
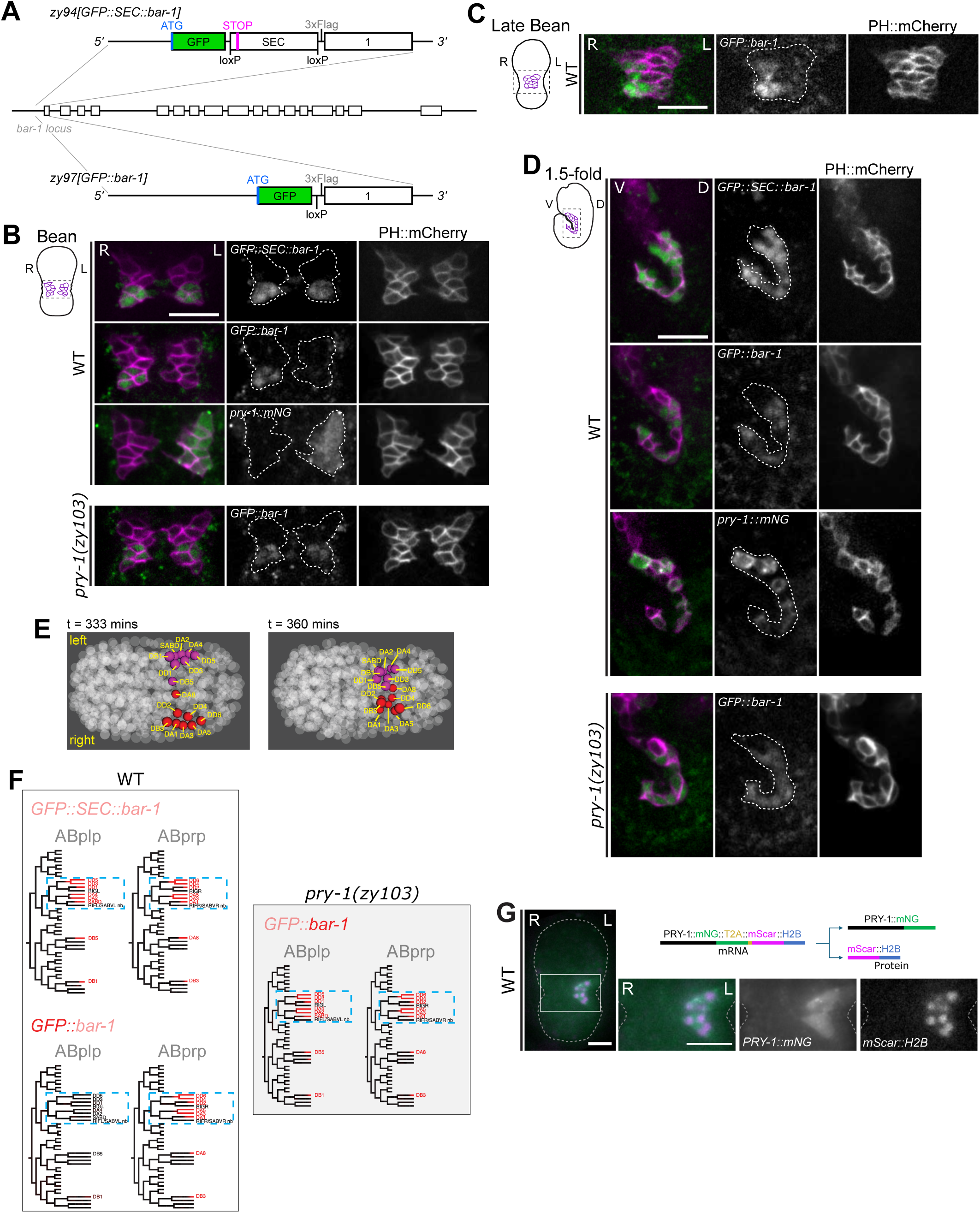
BAR-1 and PRY-1 are asymmetrically expressed on opposite sides during VNC assembly. (A) Schematic showing the locations of GFP knock-ins for endogenous transcriptional (*zy94[GFP::SEC::bar-1])* and translational (*zy97[GFP::bar-1]) bar-1* reporters. (B-D) Representative images of *zy94[GFP::SEC::bar-1]*, and *zy97[GFP::bar-1]*, in WT and *pry-1(zy103)* embryos, and *cp383[pry-1::mNG]*, an endogenous PRY-1 translational reporter, in WT embryos at bean (B), late bean (C) and 1.5-fold (D) stages. The *cnd-1p::PH::mCherry* reporter was used to label the membranes of a subset of VNC neurons in (B-D). In WT embryos, *bar-1* transcriptional activity (*zy94*) is detected on both the left and right sides (n > 30), whereas BAR-1 protein (*zy97*) is restricted to the right side (n > 30). Loss of *pry-1* results in BAR-1 protein expression on both the left and right sides (n > 30). Schematics in the top-left corner of each panel show the embryonic stage and area (dotted box) depicted in the fluorescent images. Scale bars, 10 µm. (E) 3D reference projections of bean-stage embryos showing the positions of relevant ABpl/rp neurons at 333 and 360 min post-fertilization, with expressing nuclei color-coded by lineage origin: right (red) or left (magenta). (F) Expression in ABplp and ABprp lineages from *bar-1* transcriptional and translational reporters in WT and, for the translational reporter only in *pry-1(zy103)* mutants, as measured by confocal microscopy and automated lineage tracing (StarryNite). (G) Reporter schematic and representative image of an endogenous *zy147[pry-1::mNG::T2A::mScarlet-I::H2B*cp383]* reporter showing left-sided PRY-1::mNG (green cytoplasm) and mScarlet-I::H2B (magenta nuclei) expression in bean-stage embryos, consistent with asymmetric regulation of *pry-1* expression at the transcriptional or post-transcriptional level.

To more precisely define the spatiotemporal pattern of BAR-1 expression, we imaged embryos expressing either the *bar-1* transcriptional (*zy94*) or translational (*zy97*) fusion together with a ubiquitously expressed mCherry-tagged histone marker to label nuclei. Nuclear positions and cell lineages were determined using StarryNite (Bao et al., 2006), allowing quantification of nuclear GFP fluorescence during VNC assembly (Fig. 3E and 3F). Lineage tracing at the bean stage confirmed that the *bar-1* transcriptional fusion was expressed on both the left and right sides in DD1-6 and DA1-5, including the parental cells that give rise to DD4/6 and DA3/5, as well as DB1, DB3, DB5 and DA8 neurons (Fig. 3E and 3F). Expression in the latter four neurons showed a delay and would therefore have been difficult to assign to specific neurons at these stages without lineage-based cell tracking. This same approach confirmed that the GFP::BAR-1 protein was restricted to right side-derived neurons, including DA8 and DB3. Taken together, these findings demonstrate that *bar-1* transcription occurs symmetrically in both left and right-side-derived neurons, indicating that BAR-1 asymmetry is established through a post-transcriptional mechanism.

### PRY-1 negatively regulates BAR-1 in left-side neurons during VNC assembly

*pry-1* acts as a negative regulator of *bar-1* in *C. elegans* (Gleason et al., 2002). This is consistent with our epistasis analysis, which showed that neuron position defects in *pry-1* mutants are masked in *pry-1*; *bar-1* double mutants and instead resemble those of *bar-1* mutants. To determine whether *pry-1* regulates BAR-1 levels, we examined GFP::BAR-1 expression and localization in a *pry-1* mutant background. Loss of *pry-1* resulted in ectopic BAR-1 expression in left-side neurons, producing a symmetric expression pattern beginning at the bean stage that largely mimicked the spatial and temporal *bar-1* promoter activity (Fig. 3B and 3F). Notably, GFP::BAR-1 expression on the right side did not appear earlier in *pry-1(zy103)* mutants, consistent with reduced or absent functional PRY-1 activity on the left side being insufficient to alter the timing of BAR-1 accumulation on the right. Similarly, additional BAR-1 expressing VNC cells (presumably corresponding to the left-side neurons) are observed in 1.5-fold *pry-1* embryos (Fig. 3D). This finding is consistent with our genetic evidence and provides molecular confirmation that *pry-1* negatively regulates BAR-1 expression.

This observation prompted us to investigate the expression pattern of PRY-1 during VNC assembly. To do this, we examined PRY-1 expression using the endogenously-labeled *pry-1(cp383[pry-1::mNG])* (Heppert et al., 2018) in combination with the *zyIs36[cnd-1p::PH::mCherry]* marker. Like BAR-1, we observed a strikingly asymmetric expression of PRY-1 in WT embryos, beginning at the early bean stage (Fig. 3B and 3D). In this case, however, it was localized exclusively to left-side neurons. Whereas BAR-1 expression was initially restricted to the posterior, PRY-1 localization encompassed the entire left-side array of *cnd-*1-positive neurons from the onset of expression. Although PRY-1 is expressed on the left-side, it was unclear whether this asymmetry was regulated at the transcriptional or translational level. To test this, we used CRISPR/Cas9 to insert a T2A self-cleaving peptide and mScarlet-I downstream of mNG in *pry-1(cp383)* to produce *zy147[pry-1::mNG::T2A::mScarlet-I::H2B]*. We found that *mScarlet-I::H2B* expression was also left-side asymmetric in all embryos examined (>10), consistent with a left-sided *pry-1* transcription pattern (Fig. 3G). These results indicate that PRY-1, as an Axin homolog, establishes BAR-1 asymmetry by preventing BAR-1 accumulation in left-side neurons, consistent with its canonical role in promoting β-catenin degradation.

### Notch signaling establishes PRY-1 asymmetry to break LR symmetry in BAR-1 expression

Genetic epistasis analysis using the *mig-14(ga62)* reduction-of-function allele revealed that Wnts do not appear to act upstream of BAR-1 signaling (Fig 1B and 1E). To further test this possibility, we examined whether the *mig-14* mutation affects the asymmetric expression of BAR-1. We found that *mig-14* does not alter BAR-1 expression (Fig S2), consistent with our phenotypic analysis. Thus, an alternative non-Wnt-dependent pathway may be involved in establishing BAR-1 asymmetry.

The finding that *pry-1* asymmetry contributes to breaking LR symmetry in *bar-1* expression is reminiscent of the role of Notch signaling in LR patterning during *C. elegans* embryogenesis (Hutter and Schnabel, 1995). Notch signaling is a highly conserved pathway that mediates signal transduction between neighbouring cells and is most commonly associated with specifying cell fate during animal development (Siebel and Lendahl, 2017). Signaling is initiated when a membrane-bound ligand from the DSL (Delta/Serrate/Lag-2) family expressed on one cell, binds and activates the Notch receptor on a neighboring cell, triggering receptor cleavage and release of the Notch intracellular domain (NICD), which translocates to the nucleus to activate target gene expression together with the DNA-binding protein CSL (Siebel and Lendahl, 2017). In *C. elegans*, the core components of this pathway include the DSL-family ligands LAG-2 and APX-1, the Notch receptors LIN-12 and GLP-1, and the CSL homolog LAG-1 (Lv et al., 2024).

In *C. elegans*, nearly all neurons descend from the AB founder cell, with the posterior descendants of ABpl and ABpr giving rise to left- and right-side VNC neurons respectively (J. E. Sulston et al., 1983). Notch signaling from an MS lineage descendant that contacts cells in the left, but not right, ABplp lineage is responsible for several other LR asymmetries in the AB lineage (Hutter and Schnabel, 1995; Moskowitz and Rothman, 1996; Neves and Priess, 2005). To test the role of Notch in breaking symmetry in the lineages that give rise to VNC motor neurons, we first measured nuclear mNG levels using the knock-in allele *lin-12(ljf31)*, which expresses a LIN-12::mNG fusion protein tagged at the C-terminus (Medwig-Kinney et al., 2022; Pani et al., 2022). Because ligand binding leads to Notch receptor cleavage and translocation of the C-terminally tagged NICD to the nucleus, where it interacts with CSL, nuclear mNG signal in this strain serves as a proxy for active Notch signaling. We observed higher nuclear LIN-12::mNG levels in the left ABplpp lineage (Fig. S3), as well as other left-side lineages known to have active LIN-12/Notch signaling such as ABplpap (Sarov et al., 2012), supporting the conclusion that Notch signaling is asymmetrically active in VNC neurons born on the left side.

To determine if LIN-12/Notch signaling was required to promote *bar-1* asymmetry in right-side neurons, we examined mutants in the LIN-12/Notch pathway for alterations in GFP::BAR-1 expression. We first examined *lin-12*/Notch and *lag-1*/CSL mutants, as both genes were identified in our genetic screen for DD motor neuron position defects (Table S3). This is the same screen described above that identified the *pry-1(zy103)* allele. We also examined a *lag-2*/Delta mutant, as LAG-2 is the Notch ligand expressed in MS descendants that activates LIN-12 in left-side AB lineages (Hutter and Schnabel, 1995). Because *lin-12*, *lag-1*, and *lag-2* are essential genes, all the alleles used were hypomorphic, resulting in partial loss-of-function phenotypes.

We found that disruption of each Notch pathway component resulted in fully penetrant, symmetric GFP::BAR-1 expression in both left and right-side neurons (n ≥ 6 embryos examined/genotype) (Fig. 4A). Because this phenotype resembled that of *pry-1* mutants, we also examined PRY-1::mNG expression and found that it was completely abolished in these embryos (n ≥ 6) (Fig. 4B). This loss of signal was not due to loss of the endogenous reporter background itself, as confirmed by diagnostic PCR. These findings support a model in which the ABplp lineage Notch signaling pathway induces the expression of the β-catenin destruction complex component PRY-1/Axin specifically in left-side neurons, thereby breaking BAR-1 symmetry, presumably by destabilizing BAR-1 on the left-side (Fig. 4C). Interestingly, these findings suggest that assembly of the β-catenin destruction complex may be controlled by Notch-mediated regulation of Axin expression, rather than through the post-translational mechanisms that regulate destruction complex activity in canonical Wnt signaling.

**Figure 4.**
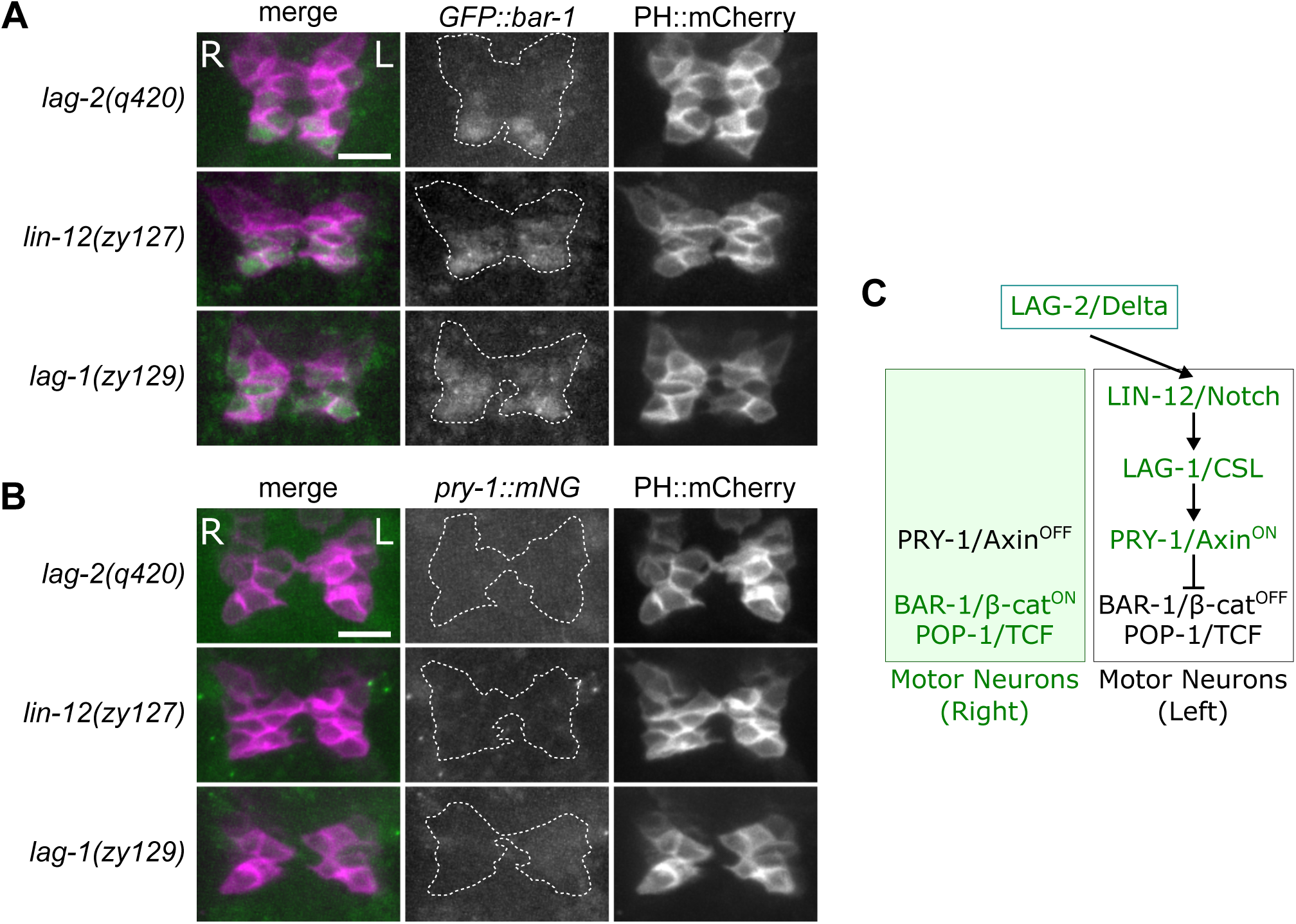
Notch signaling regulates PRY-1 to establish asymmetric BAR-1 expression. (A-B) Representative images of bean-stage embryos expressing endogenous (A) *zy97[GFP::bar-1]* or (B) *cp383[pry-1::mNG]* reporters in WT and *lag-2/delta, lin-12/notch*, and *lag-1/CSL* mutant backgrounds. *cnd-1p::PH::mCherry* was used to label a subset of VNC neuron membranes. In Notch pathway mutants *lag-2(q420)* (n = 6), *lin-12(zy127)* (n = 6), and *lag-1(zy129)* (n = 11), BAR-1 protein expression is detected on both sides of the VNC. By contrast, PRY-1 expression is abolished in *lag-2(q420)* (n = 9), *lin-12(zy127)* (n = 6), and *lag-1(zy129)* (n = 6). Scale bar, 10 µm. (C) Model showing how left-sided Notch signaling promotes PRY-1 expression in left-side neurons of the developing VNC, thereby restricting BAR-1 protein expression to the right side.

### BAR-1 is required for DD1-DD2 separation during VNC assembly

To identify when DD positioning defects first emerge, we analyzed *bar-1* and *pry-1* mutants using the *zyIs40[unc-30p::GFP cnd-1p::PH::mCherry]* reporter transgene, which labels the cytoplasm of all DD neurons with GFP and the membranes of a subset of DD and DA neurons with mCherry. At the 1.5-fold stage in WT embryos, DD1 and DD2 were in contact with one another, as were DD5 and DD6, while DD2 through DD5 were each separated by at least one intervening cell body (Fig. 5A). By the 2-fold stage, DD1 and DD2 had separated by approximately one cell body width. In *bar-1* mutants, DD1 and DD2 remained significantly closer together than in WT at the 2-fold stage (*p* = 0.0049), suggesting a failure to separate normally (Fig. 5A and 5B). In *pry-*1 mutants, DD1 and DD2 also appeared more closely positioned than in WT, although this difference did not meet statistical significance (*p =* 0.0558). These findings suggest that the *bar-1*-dependent DD1-DD2 separation defect emerges between the 1.5-fold and 2-fold stages, a period when motor neurons are establishing their AP positions along the developing VNC.

**Figure 5.**
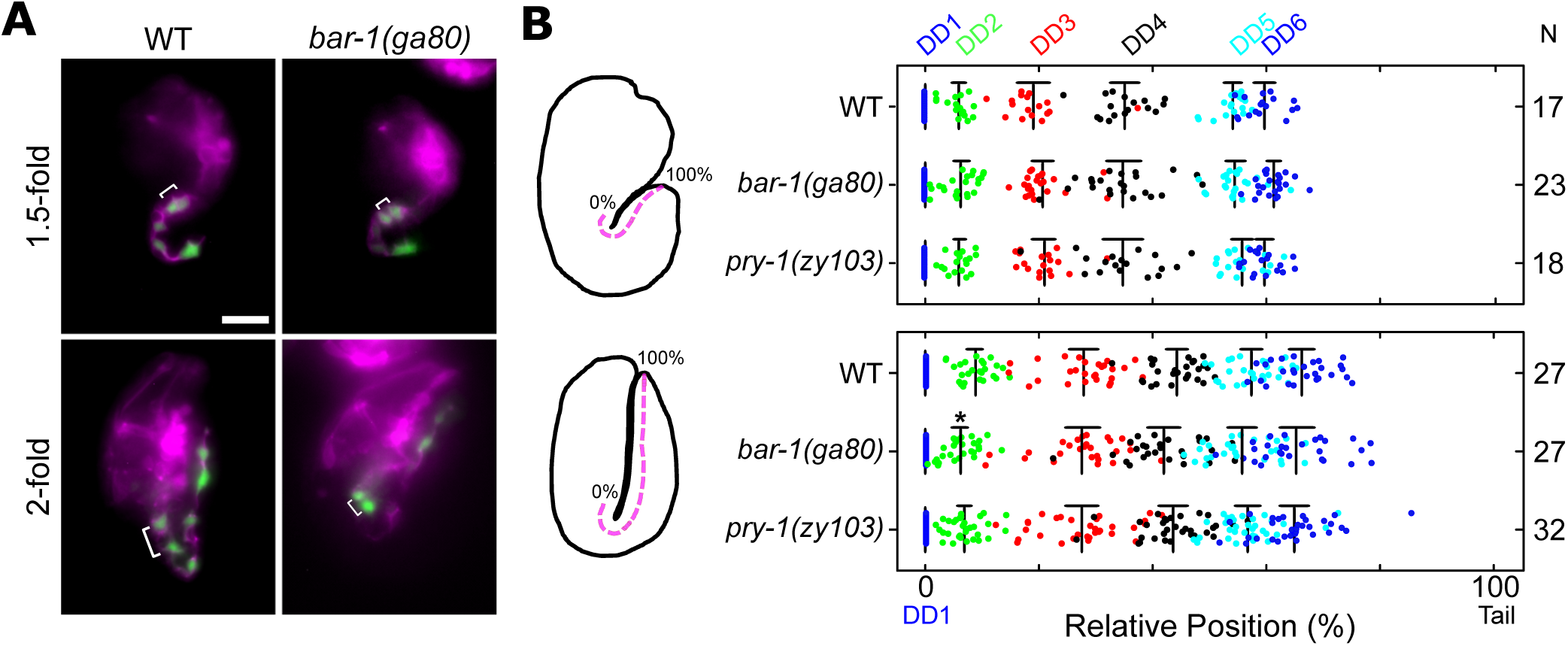
*bar-1* regulates DD1-DD2 spacing between the 1.5 and 2-fold stage of VNC assembly. (A) Representative images of WT and *bar-1(ga80)* embryos at 1.5-fold and 2-fold stages. DD neurons (green) and the VNC (magenta) are labeled with *zyIs40[unc-30p::GFP + cnd-1p::PH::mCherry]*. Brackets indicate the distance between DD1 and DD2. (B) Quantification of DD2-6 neuron positioning relative to DD1 (0%) and tail (100%) (dashed line in schematic) at 1.5-fold (top) and 2-fold (bottom) stage. *bar-1* mutants exhibit shorter DD1-DD2 spacing at 2-fold stages compared to WT. Sample sizes are indicated to the right of each plot. A one-way ANOVA was performed for each DD neuron at each stage with Dunnet’s post-hoc test when appropriate (**\***p<0.05). Scale bars, 10 µm.

### DD1-DD2 spacing defects in *bar-1* mutants arise from abnormal DA2 intercalation

During development, DD and DA neurons are born on the left and right sides of the developing VNC. As development proceeds, left-side neurons (e.g. DD1 and DA2) and their right-side counterparts (e.g. DD2 and DA3) migrate medially and form intercellular junctions with one another (Shah et al., 2017). These junctions eventually dissociate through intercalations that organize the DA, DB and DD neurons into the stereotyped L1 motor neuron pattern. One distinguishing feature of the *bar-1(ga80)* phenotype is that DD1 and DD2 were positioned closer together in both L1 larvae and 2-fold embryos (Fig. 1 and 5). Several scenarios could explain this observation: (1) DD1 and DD2 intercalate into a single file, but DD2 misintercalates anterior to DD1; (2) DD1 and DD2 intercalate into a single file, but neurons that usually intervene between them (e.g. DB3 and DA3) instead misintercalate anterior to DD1 or posterior to DD2; (3) DD1 and DD2 are unable to intercalate, leading to delays or stalling. To determine which scenario was most likely, we leveraged the asymmetric expression of *zy147[pry-1::mNG::T2A::mScarlet-I::H2B]*, described above, together with the DD nuclear marker *zySi10[unc-30p::mNG::H2B]*. This reporter combination allowed us to distinguish right-side DD neurons (DD2, DD4, DD6; *zySi10* only), left-side DD neurons (DD1, DD3, DD5; *zySi10* and *zy147* positive) and left-side DA neurons (DA2, DA4; *zy147*-only) based on their differential fluorophore expression. We found that DD1 and DD2 did not switch their relative positions in 2-fold stage *bar-1(ga80)* mutant embryos (Fig. S4A and S4B). In contrast, DD2 and DA2 switched their relative positions in some *bar-1* mutants. In WT embryos, DA2 had not yet intercalated into a single file with DD2 in 33.3% (4/12) of embryos. In the remaining 66.7% (8/12) of the embryos, DA2 intercalated exclusively anterior to DD2. In *bar-1(ga80)* mutants, DA2 remained unintercalated in 40.0% (4/10), intercalated properly anterior to DD2 in 10% (1/10), and misintercalated posterior to DD2 in 50% (5/10) of the embryos. These findings raise the possibility that the anterior displacement of DD2 observed at L1 (Fig. 1A and B) and the 2-fold (Fig. 5A and B) results from defective DD2 and DA2 intercalation at the 2-fold stage.

### Asymmetric BAR-1 expression is required for intercalation-mediated DD1-DD2 separation

We have shown that DD1-DD2 separation occurs between the 1.5-fold and 2-fold stages in WT embryos through a mechanism involving DA neurons. However, the specific intercalation events that establish the order of motor neurons observed at L1 were still unknown. Consequently, it also remained unclear how perturbing BAR-1 asymmetry affected the broader intercalation process. The role of DB neurons in VNC development is also unknown but is worth considering as DB1 and DB3 express BAR-1 (Fig. 3F) and are in direct contact with other DA and DD neurons during this period. Furthermore, *bar-1(ga80)* and *pry-1(zy103)* mutants exhibit significant albeit lower-magnitude position changes in DB1 and DB3 (Fig. 1B; Table S1).

To directly observe the cellular movements underlying VNC assembly, we used the extrachromosomal transgenic reporter *zyEx92[ceh-6p::mNG::PH + cnd-1p::mScarlet-I::PH + odr-1p::RFP]*, which labels DB neuron membranes with mNG and DA and DD neuron membranes with mScarlet-I (Fig. S5). We combined *zyEx92* and *zySi10* and performed lightsheet time-lapse microscopy to manually track the position and movement of specific DA, DB, and DD motor neurons from bean stage to beyond the 1.5-fold stage, when the embryo begins to twitch. Each embryo analyzed by this method is depicted in Fig. 6A). Using this live imaging approach, we could visualize the positioning of DB neurons between the left and right groups of DA and DD neurons. As the left and right-side DA and DD neurons converge toward the ventral midline, the DB neurons are displaced ventrally, resulting in a ventral layer of DB neurons and a dorsal layer of DA and DD neurons (Fig. S5 and Video 1). From the 1.5-fold to roughly 2-fold stage, the interconnected DB neurons separate from one another, allowing them to intercalate with cells in the dorsal DA/DD layer (Fig. S5 and Video 2).

**Figure 6.**
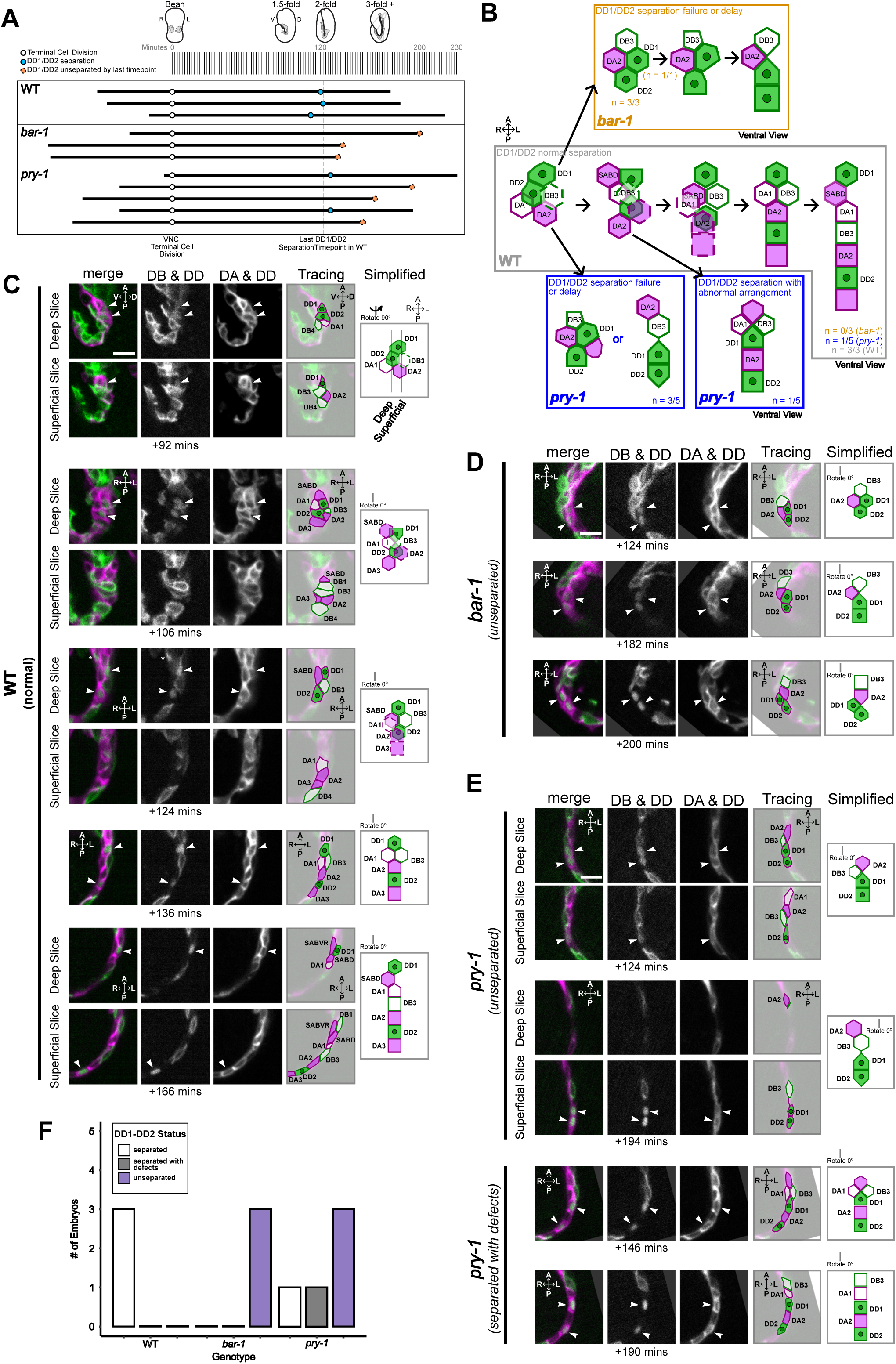
Intercalation-mediated separation of DD1 and DD2 during VNC assembly is disrupted in *bar-1* and *pry-1* mutant embryos. (A) Overview of the timelapse sample set acquired using light sheet microscopy. Each horizontal black line corresponds to individual WT, *bar-1* or *pry-1* embryos. The length of each line indicates the developmental time window covered by the corresponding timelapse. Individual timelapses were aligned at the time point at which the terminal cell divisions producing DA, DD and DB neurons were completed (+0 min). (B) Schematic summary of the WT intercalation trajectory leading to the separation of DD1 and DD2, and the corresponding phenotypes in *bar-1* and *pry-1* mutants. (C-E) Example fluorescence images, tracings, and simplified illustrations of VNC cell configurations at select time points depicting the WT trajectory (C), unseparated phenotype in *pry-1* or *bar-1* mutants (D and E) or separated with defects phenotype in *pry-1* mutants (E, lower). Motor neurons were labeled using *zyEx92[ceh-6p::mNG::PH + cnd-1p::mScarlet-I::PH]; zySi10[unc-30p::mNG::H2B]* to visualize DA (magenta membrane), DB (green membrane) and DD (magenta membrane with green nuclei) neurons. Scale bars, 5 µm. For clarity, simplified illustrations include only neurons that are relevant to DD1 and DD2 separation (SABD, DD1, DA1, DB3, DA2, DD2 and DA3). Neuron identities were assigned based on reporter expression and on the assumption that the relative order of neurons within each class (DA, DB or DD) was maintained. Additionally, a rotation maker is included to highlight if the tracings had to be rotated in 3D to orient all simplified illustrations into A-P running vertically and R-L running horizontally. (F) Quantification of the penetrance of DD1-DD2 separation phenotypes (separated, separated with defects, or unseparated) in WT, *bar-1* and *pry-1* embryos.

Since the primary defect in *bar-1(ga80)* mutants involves the inability to separate DD1 from DD2 (Fig. 5), we used this live imaging dataset to focus specifically on the cellular events underlying this separation. Tracking individual neurons over time, we observed that DD1 and DD2 begin to separate after 1.5-fold stage (Fig. 6B and 6C) and are fully separated by DA1, DB3 and DA2 by the 3-fold stage (n = 3). Time-lapse tracking revealed that this separation occurs through a series of intercalation steps. First, the formation of a two- or three-way junction involving DA1, SABD, and DB3 inserts these neurons between DD1 and DD2 to separate them. Next, DA2 intercalates anterior to DD2 and posterior to the DA1/SABD/DB3 group. Lastly, DA1 separates from DA2 to insert between DD1 and DB3. The final position of SABD varied between embryos. Overall, our time-lapse observations show that the stepwise intercalation of DA and DB cells initiates the separation of DD1 and DD2 beginning at the 1.5-fold stage.

This live imaging approach also allowed us to examine whether DD1-DD2 separation was altered by perturbing asymmetric BAR-1 expression. In both the *bar-1(ga80)* (n = 3/3) and *pry-1(zy103)* (n = 3/5) mutants (corresponding to symmetric OFF and symmetric ON BAR-1 expression states), we observed that DD1 and DD2 remained together at the developmental stage when they had separated in WT embryos (Fig. 6A-F). For *pry-1* embryos, one embryo completed DD1-DD2 separation normally (n = 1/5) and one embryo was able to separate DD1-DD2 but with an abnormal arrangement of neurons (n = 1/5). These observations are consistent with our finding that DA2 is frequently mispositioned posterior to DD2 (Fig. S4) and that DD2 is closer to DD1 (Fig. 5) in 2-fold stage *bar-1* mutant embryos. Furthermore, the variation in phenotypes observed in our *pry-1* timelapse analysis also reflects the milder DD2 positioning phenotype observed at 2-fold stage (Fig. 5). Thus, although these timelapse observations were not sufficiently powered for statistical analysis of separation dynamics, they suggest that disrupting asymmetric BAR-1 expression is associated with a failure in the initiation of DD1-DD2 separation which can contribute to the disorder of neurons.

### BAR-1 acts instructively to promote VNC motor neuron placement

Our earlier phenotypic analysis showed that reduced β-catenin signaling output, as in *bar-1* and *pop-1* mutants, causes anterior shifts in motor neuron position, whereas increased β-catenin signaling output, as in *pry-1* mutants, causes posterior shifts. The motor neurons most strongly affected by reduced β-catenin signaling were the right-side-derived neurons that express *bar-1*. For example, DD2 and DA3, which exhibited some of the strongest defects, were displaced anteriorly in *bar-1* mutants (Fig. 1B and 1C). These observations suggested that BAR-1 may provide an instructive signal that initiates movement toward their final posterior positions, including the initiation of DD1-DD2 separation observed in our time-lapse studies. To test whether BAR-1 could instruct neurons to shift posteriorly, we sought to ectopically express *bar-1* in left-side neurons that normally do not express BAR-1. Because we lacked a left-side-specific promoter, we considered using the *bar-1* promoter itself, which, unlike BAR-1 protein, is active in both left and right-side motor neurons. To determine the promoter region required to drive *bar-1* expression in VNC motor neurons, we took advantage of the detailed promoter dissection performed by Natarajan et al. (2004). They identified a 321-bp element located approximately 5 kb upstream of the *bar-1* start ATG that was sufficient to drive expression in VNC neurons. They also identified a regulatory region required for expression in the 12 Pn.p hypodermal cells, which are initially arranged along the ventral midline adjacent to the VNC. Because we wanted to restrict ectopic BAR-1 expression to neurons and avoid potential effects of BAR-1 expression in the surrounding hypodermis, we tested a *bar-1* promoter extending approximately 7 kb upstream of the start codon but containing a ∼1.9-kb deletion that removed the Pn.p regulatory region (Fig. S6A). The resulting mNG reporter line (*zyIs58*) recapitulated the embryonic VNC expression pattern observed with our endogenous *bar-1* GFP knock-in reporter (*zy94*) (Fig. S6B).

We next asked whether the level of BAR-1 expression influences VNC neuron positioning. In *pry-1* mutants, BAR-1 is ectopically expressed on the left side (Fig. 3) and motor neurons are mispositioned towards the posterior (Fig. 1), suggesting that increased BAR-1 signaling may contribute to the posterior displacement of neurons. To increase BAR-1 expression further, we generated transgenic animals carrying a multicopy integrated array (*zyIs59[bar-1p::BAR-1]*) in which the modified *bar-1* promoter in *zyIs58* was fused to a *bar-1* cDNA to drive BAR-1 expression in both left and right-side DD and DA neurons. *zyIs59* alone did not significantly alter the mean position of VNC neurons, possibly because endogenous PRY-1 remained present and continued to suppress ectopic BAR-1 protein in left-side neurons (Fig. 7A). We then crossed *zyIs59* into the *pry-1* mutant background, thereby increasing BAR-1 expression in the absence of its negative regulator. We found that several neurons which were mispositioned in *pry-1* mutants alone were displaced further posteriorly in *pry-1; zyIs59* animals. Other neurons, including DD3, DD4, and DD5, which were positioned normally in *pry-1* mutants, displayed severe posterior shifts in the presence of *zyIs59* (Fig. 7A and 7B). The pronounced posterior displacement of DD3-DD5 was particularly striking, as these mid to posterior VNC neurons appeared substantially more sensitive to ectopic BAR-1 than the DA and DB neurons located in the posterior VNC. This differential response suggests that DD neurons may be intrinsically more responsive to BAR-1, but that their response is normally constrained by mechanisms that limit BAR-1 activity or downstream signaling. Overexpression of *bar-1* in the *pry-1* mutant background may overcome these constraints, revealing an underlying capacity of DD neurons to respond to BAR-1. Overall, these findings indicate that BAR-1 levels influence VNC motor neuron positioning and support a model in which BAR-1 acts instructively to promote posterior displacement.

**Figure 7.**
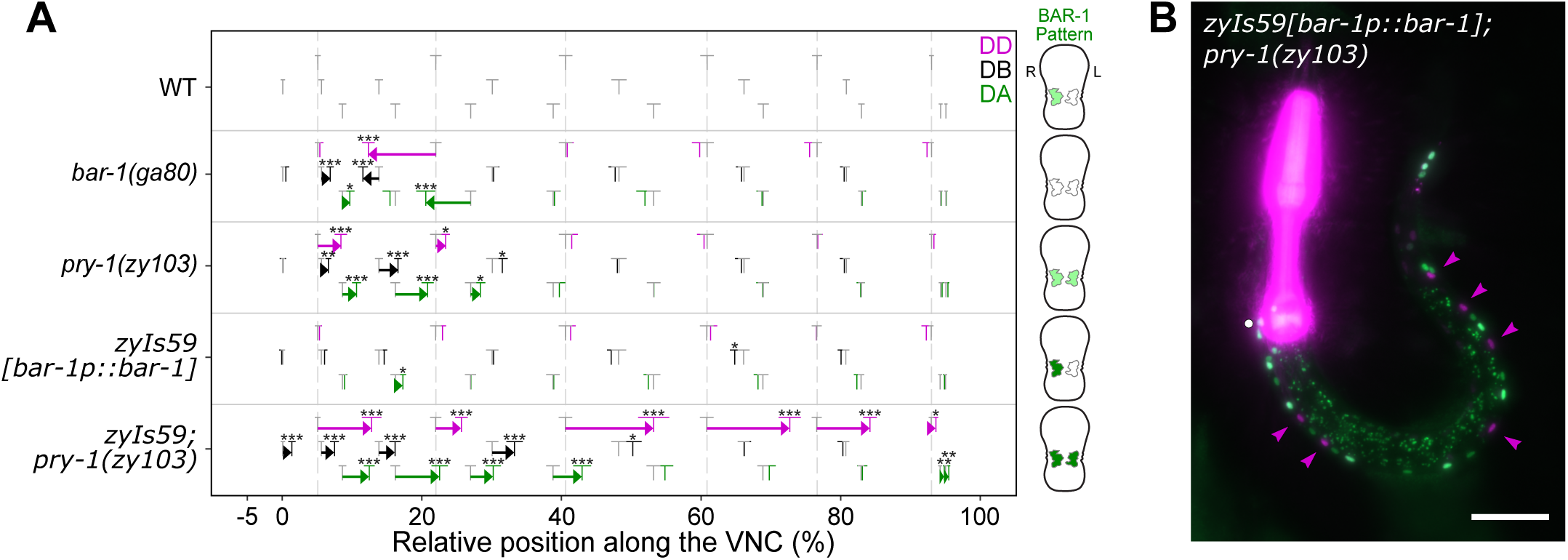
BAR-1 acts instructively to promote posterior positioning of motor neurons. (A) Plot of relative mean position of DD, DA, and DB motor neurons in WT, mutants, and *bar-1* overexpression strains: *zyIs59[bar-1p::BAR-1]* and *zyIs59; pry-1(zy103)*. For ease of comparison with the overexpression strains, the WT, *bar-1(ga80),* and *pry-1(zy103)* datasets from Figure 1B were replotted. When *bar-1* is overexpressed in the *pry-1(zy103)* background, motor neurons exhibit strong posterior shifts, with the largest shifts observed in DD neurons. Embryo schematics indicate the left-right asymmetry and relative BAR-1 expression levels associated with each genotype. Higher expression levels are represented as a darker green colour. Sample sizes as well as other statistical values are listed in Table S1. Mutant positions were compared to WT using Welch’s *t*-test followed by multiple comparisons correction using the Benjamini-Hochberg procedure for controlling the false discovery rate to obtain adjusted *p* values (performed alongside comparisons in Figure 1B). Annotations as in Fig 1B. (B) Representative fluorescence image of *zyIs59; pry-1(zy103)* with motor neurons labeled as in Fig. 1A. Magenta arrowheads indicate posteriorly displaced DD neurons. The pharynx is labeled with the *myo-2p::mCherry* (magenta) co-injection marker used to generate *zyIs59*. Scale bar, 100 µm.

### BAR-1 activity modulates POPTOP expression in the developing VNC

BAR-1 is the *C. elegans* β-catenin paralog most closely associated with canonical Wnt transcriptional regulation. It does so by binding the DNA binding transcription factor POP-1 to regulate target gene transcription (Korswagen et al., 2000). Our earlier findings that BAR-1 is expressed in the nucleus throughout VNC morphogenesis (Fig. 3B-D) and that *pop-1(hu9)* mutants exhibited DD1 and DD2 positional defects similar to those of *bar-1* null mutants, while *pop-1; bar-1* double mutants displayed a phenotype of similar severity to *bar-1* single mutants (Fig. 1B and 1D) prompted us to further investigate the relationship between BAR-1 and POP-1 during VNC assembly. To this end, we used POPTOP, a transcriptional reporter consisting of seven POP-1 binding sites positioned upstream of a minimal promoter driving mCherry (Green et al., 2008).

POPTOP expression was analyzed in combination with *zySi6[bar-1p::mNG::PH]*, which uses the same modified *bar-1* promoter utilized in our ectopic expression experiments to drive membrane-bound mNG in DD1-6 and DA1-5. In WT bean stage embryos, POPTOP expression within the VNC was predominantly higher on the right side (Fig. 8A and 8B), reminiscent of the asymmetric expression pattern observed with BAR-1 (Fig. 3B). In *bar-1* mutants, POPTOP expression appeared symmetric, with no significant difference between left and right-side VNC neurons (Fig. 8A and 8B). In addition to the loss of LR asymmetry, overall POPTOP expression within the VNC was reduced in *bar-*1 mutants (Fig. 8C). We next focused our analysis on 2-fold embryos, the stage at which VNC defects first became apparent in *bar-1(ga80)* mutants (Fig. 5 and 6). At this stage, POPTOP reporter activity was detected in the WT VNC (Fig. 8D). However, in *zyIs59; pry-1(zy103)* embryos, in which *bar-1* was overexpressed on both sides of the VNC, mean POPTOP fluorescence intensity was nearly two-fold higher than in WT embryos (Fig. 8E; *p* = 0.0135). These observations demonstrate that POP-1 transcriptional activity in the VNC depends on BAR-1, supporting a model in which the canonical POP-1/BAR-1 complex promotes target gene expression in right-side neurons to regulate VNC assembly.

**Figure 8.**
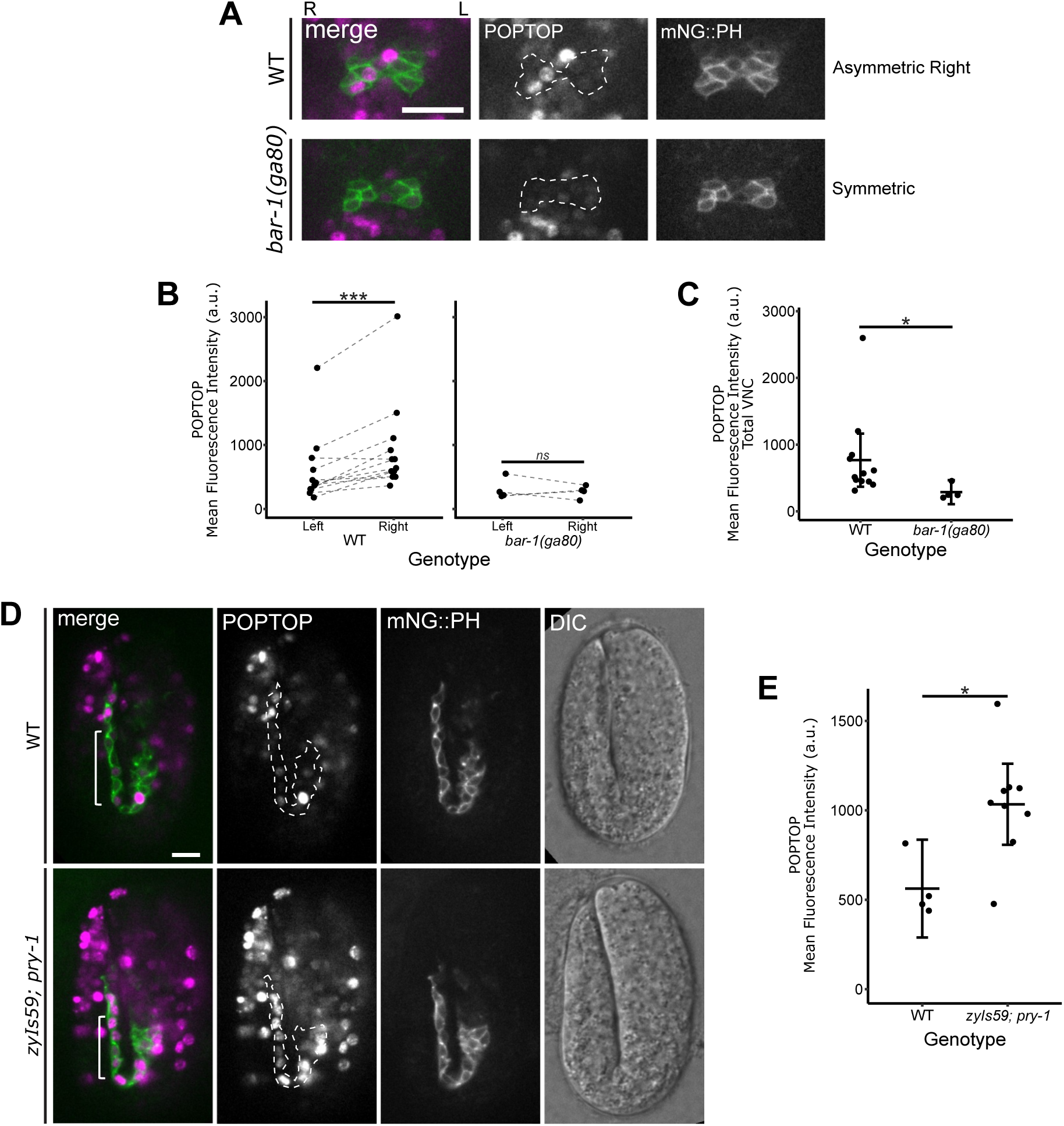
BAR-1 regulates asymmetric POP-1/TCF activity during VNC assembly. (A) Representative maximum-intensity projections of WT and *bar-1(ga80)* bean stage embryos carrying POPTOP, a reporter of POP-1/TCF-induced transcription. In WT bean-stage embryos, POPTOP expression was predominantly higher on the right side within the VNC (n = 12 embryos), whereas *bar-1* mutants displayed symmetric POPTOP expression (n = 4 embryos). Images were contrast-adjusted identically to facilitate visual comparison of fluorescence intensity. The developing VNC is labeled with membrane-localized mNG using *zySi6[bar-1p::mNG::PH]*. Scale bar, 10 µm. (B and C) Dot plots showing mean POPTOP fluorescence intensity within left versus right VNC cells (B) or within all VNC cells (C). VNC cells were segmented using the *zySi6* (mNG) channel and background fluorescence was subtracted from the measurements. Measurements were compared using Welch’s t-test (*p < 0.05, ***p < 0.001). (D) Representative maximum-intensity projections of WT and *zyIs59[bar-1p::BAR-1]; pry-1(zy103)* double mutant 2-fold embryo. Reporter expression and image adjustments were as described in (A). White brackets indicate an anatomically comparable region between strains in which differences in POPTOP expression are most apparent. Scale bar, 10 µm. (E) Dot plot showing mean POPTOP fluorescence intensity within all VNC cells at the 2-fold stage. VNC cells were segmented using the *zySi6* channel. Welch’s t-test was used to compare WT with *zyIs59; pry-1(zy103)*, *p < 0.05. Sample sizes were n = 4 (WT) and n = 9 (*zyIs59; pry-1(zy103)*).

## DISCUSSION

### β-Catenin-TCF signaling controls anteroposterior motor neuron positioning in the VNC

Here we show that BAR-1/β-catenin and its transcriptional partner POP-1/TCF regulate the AP positioning of DD, DA, and DB motor neuron cell bodies in the anterior VNC. Loss of *bar-1* or *pop-1* causes anterior shifts in neuronal cell body position, whereas loss of *pry-1*/Axin, a negative regulator of β-catenin, causes posterior shifts. We further show that Notch, PRY-1, BAR-1 and POP-1 exhibit a coordinated series of LR asymmetries in the VNC, in which Notch-dependent PRY-1 asymmetry restricts BAR1 accumulation and contributes to asymmetric POP-1 activity. Thus, our findings reveal a previously unrecognized role for an initial Notch-dependent LR asymmetry in regulating asymmetric β-catenin/TCF activity that helps determine motor neuron positioning along the AP axis. These findings identify motor neuron cell body positioning along the AP axis as an additional β-catenin-dependent neurodevelopmental process, extending the previously described roles of BAR-1 in neuronal migration, axon guidance, and synapse formation (Maloof et al., 1999; Maro et al., 2009; Tulgren et al., 2014).

The β-catenin-TCF-dependent process we identified is part of a broader, coordinated series of cell movements that assemble the VNC. We previously showed that between the bean and 1.5-fold stages of embryonic development, DD, DA, and DB neurons undergo convergent extension, directed intercalation, and neighbour-neighbour exchanges to establish the nascent VNC (Shah et al., 2017). How the VNC achieves its final stereotyped organization of motor neuron cell bodies between the 1.5-fold stage and hatching is less well understood, although it clearly involves additional sorting and positioning events. Our live-imaging data provides insight into this later phase, revealing that DD1 and DD2, which are initially positioned close together in the anterior VNC at the 1.5-fold stage, are progressively separated by the successive intercalation of other motor neurons between them. The positional defects we observe in *bar-1* and *pry-1* mutants therefore appear to reflect disruptions in this separation process.

Consistent with the idea that β-catenin activity can influence cell sorting in a different developmental context, studies in *Xenopus* have shown that the level of β-catenin activity determines the direction of cell sorting at the notochord-somite boundary, with increased activity promoting sorting towards the somites and reduced activity promoting sorting towards the notochord (Reintsch et al., 2005). Together, with our observations that BAR-1 levels can drive opposite shifts along the AP axis, these results suggest that β-catenin-TCF-activity can influence the cellular behaviours that determine cell position during tissue morphogenesis more broadly. In the VNC, our findings specifically implicate BAR-1-POP-1 activity in the intercalation and sorting behaviours that establish the final relative positions of motor neurons.

### LR-asymmetric BAR-1/POP-1 activity contributes to DD neuron separation during VNC assembly

Our analysis revealed a striking LR asymmetry in BAR-1/β-catenin expression during VNC morphogenesis. BAR-1 is expressed in right-side-derived motor neurons but is absent from their left-side counterparts. In contrast, PRY-1/Axin is asymmetrically expressed in left-side neurons, where it could restrict BAR-1 accumulation. Consistent with PRY-1 acting as a negative regulator of BAR-1, loss of *pry-1* results in BAR-1 expression on both sides of the VNC. Our live imaging showed that DD1 and DD2, which initially lie in close proximity, become increasingly separated as neighboring DA and DB neurons intercalate between them, positioning DD2 progressively more posterior to DD1. Loss of BAR-1 asymmetry in both *bar-1* and *pry-1* mutants disrupts DD1-DD2 separation, supporting a functional role for LR asymmetric BAR-1 expression in this process. Further evidence for the functional importance of this asymmetry comes from our BAR-1 overexpression experiments, in which expressing BAR-1 in all DD and DA neurons drives their cell bodies posteriorly, consistent with BAR-1 promoting posterior movement and suggesting that its asymmetric expression normally restricts this activity to right-side neurons.

The asymmetric pattern of BAR-1 expression is also reflected in asymmetric POP-1 activity, consistent with the established role of BAR-1 as a transcriptional coactivator for POP-1 (Korswagen et al., 2000). *bar-1* and *pop-1* mutants exhibit similar neuronal positioning defects. In WT embryos, we found that POPTOP expression, a reporter of POP-1 transcriptional activity, was predominantly higher on the right side, whereas this asymmetry was lost in *bar-1* mutants. Conversely, expressing BAR-1 on both sides of the VNC increased overall POPTOP activity. Together, these observations suggest that BAR-1 regulates the spatial pattern of POP-1 activity during VNC assembly, although whether this reflects a direct effect of BAR-1 on POP-1 or an indirect mechanism remains unknown. Although asymmetric POP-1 activity is well established during asymmetric cell divisions in *C. elegans*, where it differs between anterior and posterior daughter cells (Huang et al., 2007), whether POP-1 activity is also asymmetric long the LR axis has remained unclear. In zebrafish, Tcf7l2 is expressed in habenular neurons on both sides of the brain and is required for LR-asymmetric differentiation of habenular neurons, with its activity differentially regulated by signals from the left-sided parapineal (Hüsken et al., 2014). Similarly, our findings reveal LR-asymmetric POP-1-mediated transcriptional activity in the VNC that is associated with asymmetric BAR-1 accumulation.

BAR-1/POP-1 activity also influences Q neuroblast migration in *C. elegans*, where asymmetric Wnt signaling establishes distinct migratory behaviours along the AP axis: QL descendants migrate posteriorly through a canonical Wnt/β-catenin pathway, whereas QR descendants migrate anteriorly through a β-catenin independent pathway (Maloof et al., 1999; Rella et al., 2016). In the Q neuroblast lineage, canonical Wnt signaling activates BAR-1 and POP-1-dependent expression of the Hox gene *mab-5* in QL, but not QR, thereby coupling the LR difference in Wnt pathway activity to asymmetric AP migration. In the Q neuroblast lineage, PRY-1 acts through the destruction complex to negatively regulate BAR-1 stability (Korswagen et al., 2002), whereas our findings suggest that during VNC assembly, Notch signaling acts upstream of PRY-1 by regulating its expression to establish asymmetric BAR-1 accumulation.

The downstream cellular behaviors are also likely distinct. Q neuroblast migration is a cell-autonomous process in which QL migrates posteriorly, whereas DD1-DD2 separation involves the intercalation of neighboring DA and DB neurons between them. Our BAR-1 overexpression experiments, in which BAR-1 expression in both left- and right-side neurons caused severe posterior displacement of multiple VNC neurons, support the idea that BAR-1 can act cell autonomously to promote posterior movement. However, proper DD1-DD2 separation is unlikely to be entirely cell autonomous, because it requires the coordinated intercalation of neighboring neurons that do not normally express BAR-1. Our findings suggest that BAR-1 asymmetry, rather than simply BAR-1 expression, is important for proper neuronal positioning during VNC assembly. BAR-1/POP-1 transcriptional activity may therefore regulate properties of BAR-1-expressing neurons such as cell polarity, adhesion, or interactions with intercalating neighbours, thereby promoting DD1-DD2 separation. Identifying the relevant BAR-1/POP-1 transcriptional targets in the developing VNC will be necessary to distinguish among these possibilities.

### Notch-dependent regulation of Axin establishes LR asymmetry in β-catenin activity

We found that BAR-1 is asymmetrically expressed in right-side VNC neurons during assembly and that this asymmetry is established through asymmetric Notch signaling. Specifically, LAG-2/Delta activates LIN-12/Notch signaling in left-side neurons, where LIN-12 and its transcriptional effector LAG-1/CSL promote *pry-1*/Axin transcription, which in turn restricts BAR-1 accumulation to right-side neurons. In the canonical Wnt pathway, Axin acts as a scaffold within the destruction complex alongside APC and GSK-3 to promote β-catenin phosphorylation and proteasomal degradation. In this context, Axin function is regulated post-translationally through control of its stability, localization, and interactions with other destruction complex proteins (Song et al., 2014). Our findings instead suggest that Notch signaling regulates β-catenin activity through asymmetric regulation of *pry-1*/Axin transcription, revealing a mode of Axin regulation distinct from canonical Wnt signaling.

Crosstalk between the Notch and Wnt pathways has been documented in multiple developmental contexts, and several mechanisms by which Notch can suppress β-catenin activity have been described. Studies in *Drosophila* have shown that Notch can negatively regulate Armadillo/β-catenin independently of its canonical transcriptional activity by physically associating with Armadillo and promoting its sequestration within endosomal compartments, thereby limiting its nuclear accumulation and transcriptional activity (Hayward et al., 2005; Sanders et al., 2009). Subsequent work further demonstrated that components of the β-catenin destruction complex, including Axin and APC, regulate this ligand-independent trafficking of Notch, linking endosomal Notch dynamics to the availability of transcriptionally active Armadillo (Muñoz-Descalzo et al., 2011). In vertebrate cells, two distinct mechanisms have been demonstrated: at the membrane, ligand-independent endocytic trafficking that sequesters β-catenin away from TCF; and in the nucleus, an NICD-dependent mechanism that can disrupt β-catenin-induced transcription (Acar et al., 2021). In these examples, Notch limits β-catenin activity through post-translational mechanisms. In contrast, our findings point to a distinct mechanism whereby Notch regulates *pry-1*/Axin expression which in turn regulates β-catenin activity.

### LR asymmetry during nerve cord assembly organizes motor neurons along the AP axis

LR asymmetries within the nervous system have been described in a wide range of animals, from the lateralization of language areas in the human cerebral cortex to the functionally asymmetric ASE chemosensory neurons of *C. elegans* (Güntürkün et al., 2020; Johnston and Hobert, 2003). These asymmetries can establish persistent differences between the left and right sides of the mature nervous system, including differences in neuronal identity, connectivity, or function. Wnt/β-catenin signaling has also been implicated in establishing such left-right differences during nervous system development. For example, Wnt/Axin1/β-catenin signaling regulates asymmetric Nodal pathway gene expression and the lateralization of CNS structures, such as the habenulae in zebrafish (Carl et al., 2007; Hüsken and Carl, 2013).

Our results suggest that LR asymmetry can influence the AP positioning of motor neuron cell bodies during VNC assembly. This LR-to-AP coupling arises because VNC motor neurons that will ultimately occupy a single midline tract begin development as bilaterally symmetric pairs, and their final AP positions after convergence depend on the transcriptional programs that were established on the two sides before intercalation. In our system, we propose that an initial LR asymmetry in Notch signaling leads to a sequence of molecular asymmetries: Notch-dependent *pry-1* expression in left-side neurons restricts BAR-1 accumulation to right-side neurons, which in turn contributes to asymmetric POP-1 activity. Thus, AP positioning may reflect not only signals acting along the AP axis, but also molecular differences established earlier along the LR axis that influence cell intercalation and sorting behaviour and are thereby translated into AP organization during VNC assembly.

In vertebrate spinal cord development, motor neuron subtypes are organized with remarkable precision along both the AP and dorsoventral axes, an organization achieved through the coordinated action of morphogen gradients, transcriptional programs, and local cell interactions (Sagner and Briscoe, 2019). During neural tube formation, bilaterally derived neuroepithelial cells undergo convergent extension and mediolateral intercalation toward the midline (Nikolopoulou et al., 2017). Whether transient LR-asymmetric signaling contributes to these behaviours or influences the subsequent AP organization of the neural tube will require determining whether LR-asymmetric signaling occurs in these cells and whether it influences their intercalation or positioning. The *C. elegans* VNC provides a tractable system for dissecting how transient LR asymmetries can be translated into stable AP organization during neural development, using genetics and live imaging at single-cell resolutions.

## Acknowledgements

We thank David Eisenmann for providing the *bar-1* cDNA pDE248. Some strains were provided by the *Caenorhabditis* Genetics Center, which is funded by the National Institutes of Health Office of Research Infrastructure Programs (P40 OD010440). *C. elegans* allele sequence data were obtained from WormBase (http://www.wormbase.org, release WS294). The authors acknowledge the Cell Biology and Image Acquisition Core (RRID: SCR_021845) supported by the University of Ottawa, the Natural Sciences and Engineering Research Council of Canada, and the Canada Foundation for Innovation. We also acknowledge the assistance of the StemCore Laboratories Genomics Core Facility (OHRI), RRID:SCR_012601.

## Funding

This work was supported by Canadian Institutes of Health Research (CIHR) Project Grants (123513 and 156160) to A. Colavita; a Natural Sciences and Engineering Research Council of Canada (NSERC) Discovery Grant (RGPIN-2025-05223) and a Digital Research Alliance of Canada (DRAC) Resources for Research Groups grant (RRG-TPERKINS) to TJ Perkins; and National Institutes of Health (NIH) grant R35GM153497 from the National Institute of General Medical Sciences (NIGMS) to JI Murray.

**Figure S1. Reduced spacing between DD1 and DD2 motor neurons in *bar-1*, *pry-1*, and *pop-1* mutants.**

Representative fluorescent images of WT, *bar-1(ga80)*, *pry-1(mu38)*, and *pop-1(hu9)* animals expressing the *ynIs37[flp-13p::gfp]* reporter to label DD motor neurons. Brackets mark the spacing between DD1 and DD2. Compared with WT animals, DD1 and DD2 are positioned closer together in *bar-1(ga80)*, *pry-1(mu38)*, and *pop-1(hu9)* mutants. Scale bar, 20 μm.

**Figure S2. BAR-1 protein remains restricted to right-side VNC neurons in *mig-14* mutants.**

Representative image of *zy97[GFP::bar-1]* at the bean stage in a *mig-14(ga62)* mutant embryo. The *zyIs36[cnd-1p::PH::mCherry]* reporter was used to label the membranes of a subset of neurons in the developing VNC. In WT embryos, BAR-1 protein is restricted to right-side VNC neurons. This asymmetric pattern was unchanged in *mig-14* mutants (n = 8). The dotted perimeter marks the boundary of *cnd-1* promoter expression in VNC neurons. Scale bar, 10 µm.

**Figure S3. Lineage tracing of LIN-12::mNG expression.**

LIN-12::mNG nuclear expression (total nuclear intensity minus baseline/autofluorescence estimated from LIN-12(-) cells) as measured by confocal microscopy and automated lineage tracing (StarryNite) in ABpl vs ABpr lineages. Mean of n = 6 embryos. Highlighted (ABplppap and ABprppap) lineages produce several VNC neurons including DD1-6 and DA1-5.

**Figure S4. DD1 and DD2 retain their relative order in *bar-1* mutants, whereas DA2 and DD2 can switch positions.**

(A) Representative fluorescent images of WT and *bar-1(ga80)* mutant 2-fold embryos expressing the *zy147[pry-1::mNG::T2A::mScarlet-I::H2B*cp383]* reporter (see also Figure 2F) and *zySi10[unc-30p::mNG::H2B]*. This combination of reporters labels left-originating VNC cells with a magenta nuclear signal (DA and DD) and all DD neurons with a green nuclear signal allowing left DA (magenta), left DD (white), and right DD (green) to be distinguished. Diagrams beneath the images highlight the relative positions of DD1 and DD2 and of DA2 and DD2. Scale bar, 10 µm. (B) Quantification of the proportion of embryos in which DD1 is positioned anterior (normal order) or posterior to DD2 in WT and *bar-1(ga80)* mutants. (C) Quantification of the proportion of embryos in which DA2 is positioned anterior (normal order), posterior to, or unintercalated with DD2 in WT and *bar-1(ga80)* mutants. The frequencies of neuron order phenotypes between WT and *bar-1* embryos were compared using Fisher’s exact test (**p < 0.01).

**Figure S5. Visualization of DD/DA and DB neurons during VNC assembly.**

Still images from a time-lapse series showing VNC assembly in WT embryos expressing *zyEx92[cnd-1p::PH::mCherry + ceh-6p::mNG::PH]*. In the VNC, the *cnd-1p::PH::mCherry* reporter (top row) labels the membranes of DD1-6, DA1-5, and DB1 and DB3 neurons, whereas *ceh-6p::mNG::PH* (middle row) labels the membranes of predominantly DB neurons. Both reporters are also expressed in other cells. Images are shown at the indicated embryonic stages from the bean stage through the 2-fold stage, with schematic diagrams below indicating the corresponding developmental stages. During VNC assembly, neurons that form the VNC converge towards the midline and become organized into two layers by the 1.5-fold stage, with a dorsal DD/DA layer and a ventral DB layer. These neurons subsequently intercalate to form a single file by the 2-fold stage. During the time-lapse, the embryo rotates, resulting in a change in orientation from a ventral view to a lateral view. R, right; L, left; D, dorsal; V, ventral. Scale bar, 10 μm.

**Figure S6. The *bar-1* promoter used for ectopic expression.**

(A) Diagram of the *bar-1* promoter fragment used for ectopic expression analysis. The promoter extends approximately 6.8 kb upstream of the *bar-1* start codon and contains an approximately 1.9 kb deletion. The A of the ATG is designated as position 0. The promoter fragment was fused to *mNG* to generate the *zyIs58[bar-1p::mNG]* reporter. (B) Representative fluorescent images of *zyIs58; zyIs36* embryos at the bean and 1.5-fold stages, showing *bar-1* promoter activity during VNC assembly. The embryonic expression pattern driven by the *bar-1* promoter recapitulates the VNC expression pattern observed with the endogenous BAR-1 GFP knock-in allele. Schematic diagrams indicate the corresponding developmental stages. Scale bar, 10 µm.

**Supplementary Video 1. DA, DB and DD neuron movement during bean-stage development**

Timelapse video of a WT bean stage embryo expressing *zyEx92[ceh-6p::mNG::PH + cnd-1p::mScarlet-I::PH]* to label DD and DA neurons (including DB1 and DB3, but excluding DA6-DA9) with membrane-localized mScarlet-I and DB neurons (including DA8) with membrane-localized mNG. As development progresses the left and right-born mScarlet-I::PH-labelled neurons converge towards the ventral midline, while the mNG::PH-labelled neurons are displaced to a position ventral to the mScarlet-I::PH-expressing neurons.

**Supplementary Video 2. DA, DB and DD neuron movement from 1.5-fold to 2-fold development**

Continuation of the timelapse Video 1 showing a WT 1.5-fold to 2-fold stage embryo expressing *zyEx92[ceh-6p::mNG::PH + cnd-1p::mScarlet-I::PH]* to label DD and DA neurons (including DB1 and DB3, but excluding DA6-DA9) with membrane-localized mScarlet-I and DB neurons (including DA8) with membrane-localized mNG. At the start of the 1.5-fold stage, the mScarlet-I::PH-expressing neurons lie dorsal to the ventral layer of mNG::PH-expressing neurons. As development progresses to the 2-fold stage, the neurons within each layer begin to intercalate into a single file of motor neurons.

