## Supplementary figures and images for "Notch-dependent left-right asymmetry in β-catenin-TCF activity instructs anteroposterior positioning of motor neurons in the *C. elegans* ventral nerve cord"

### Supplemental Figures

**FIG S1**

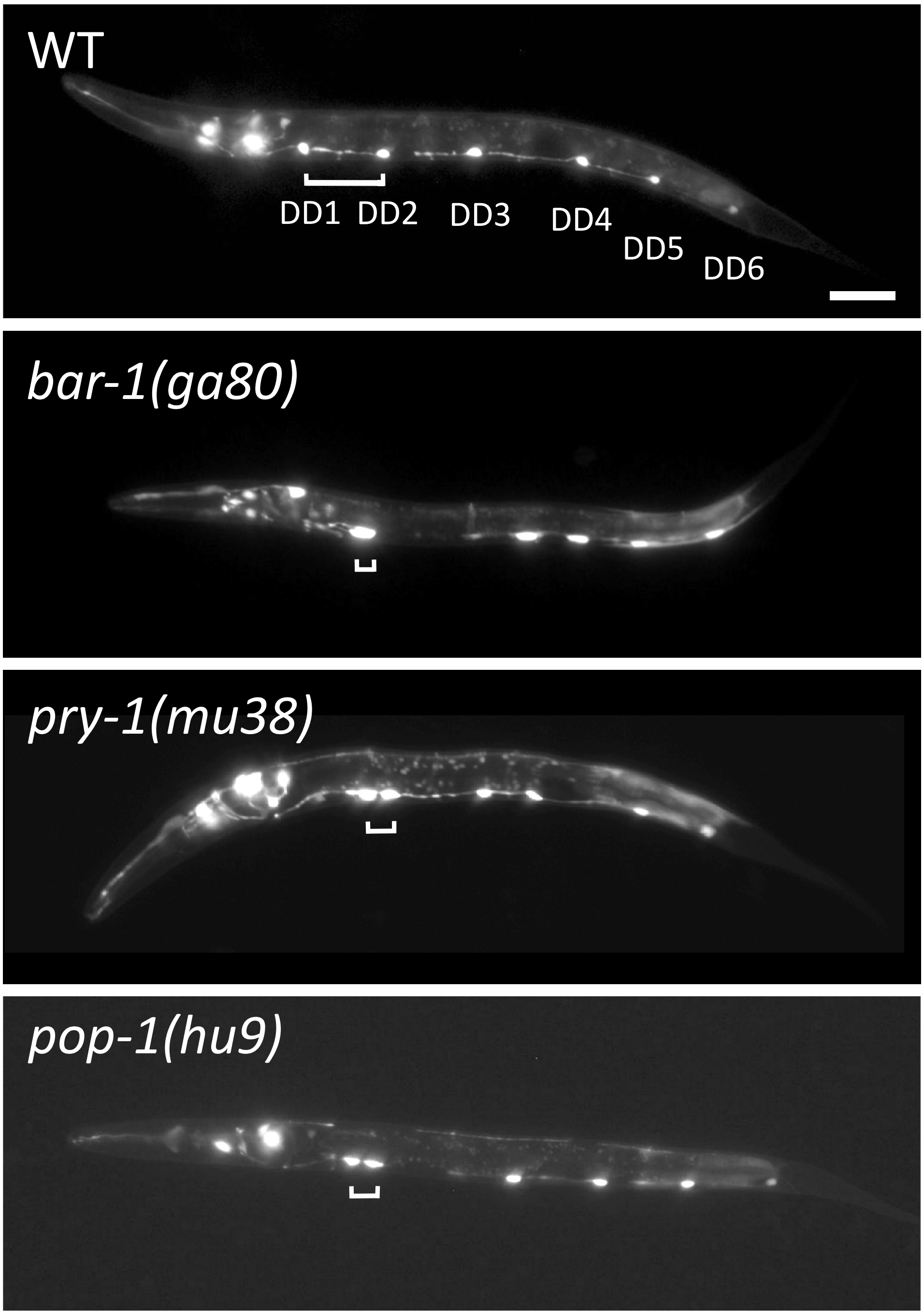

**FIG S2**

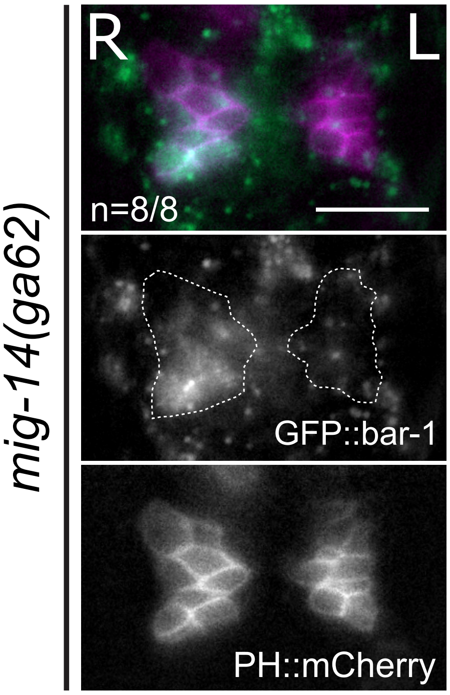

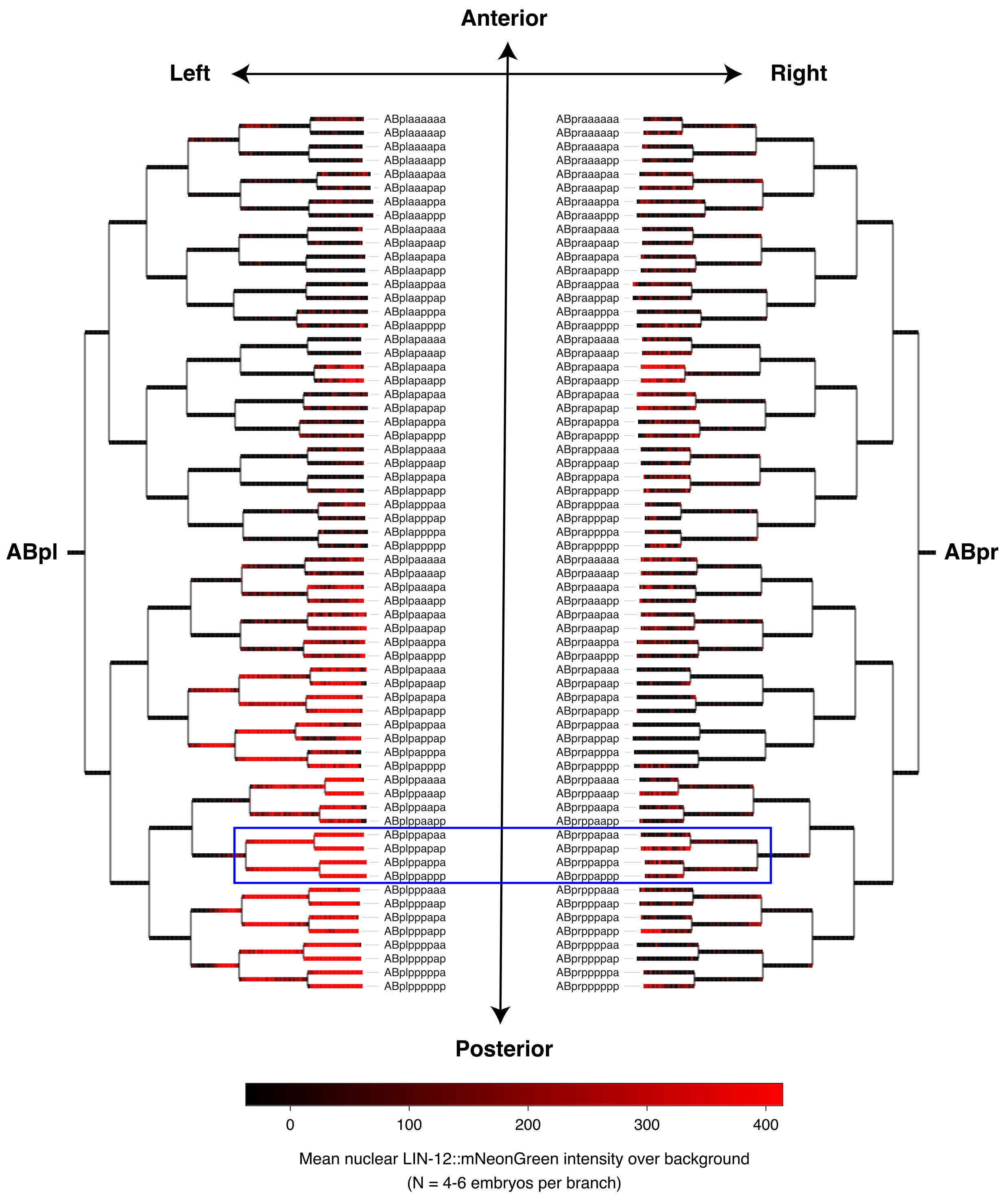

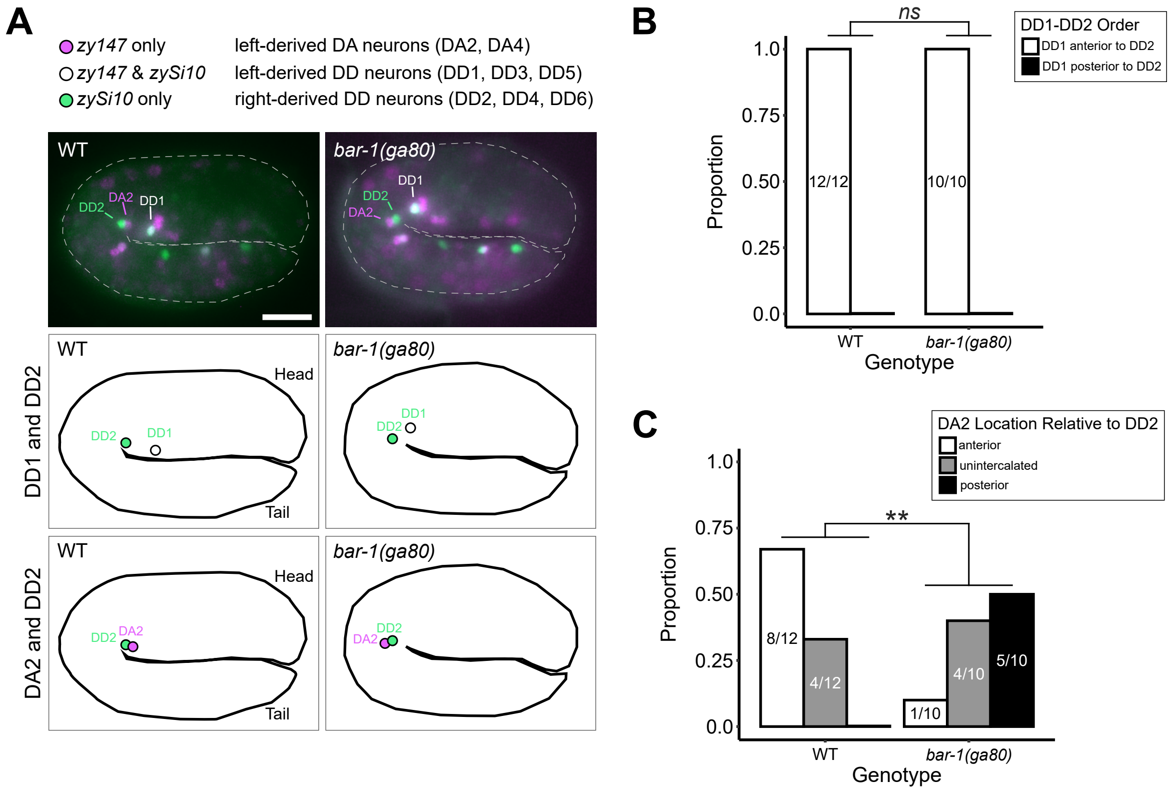

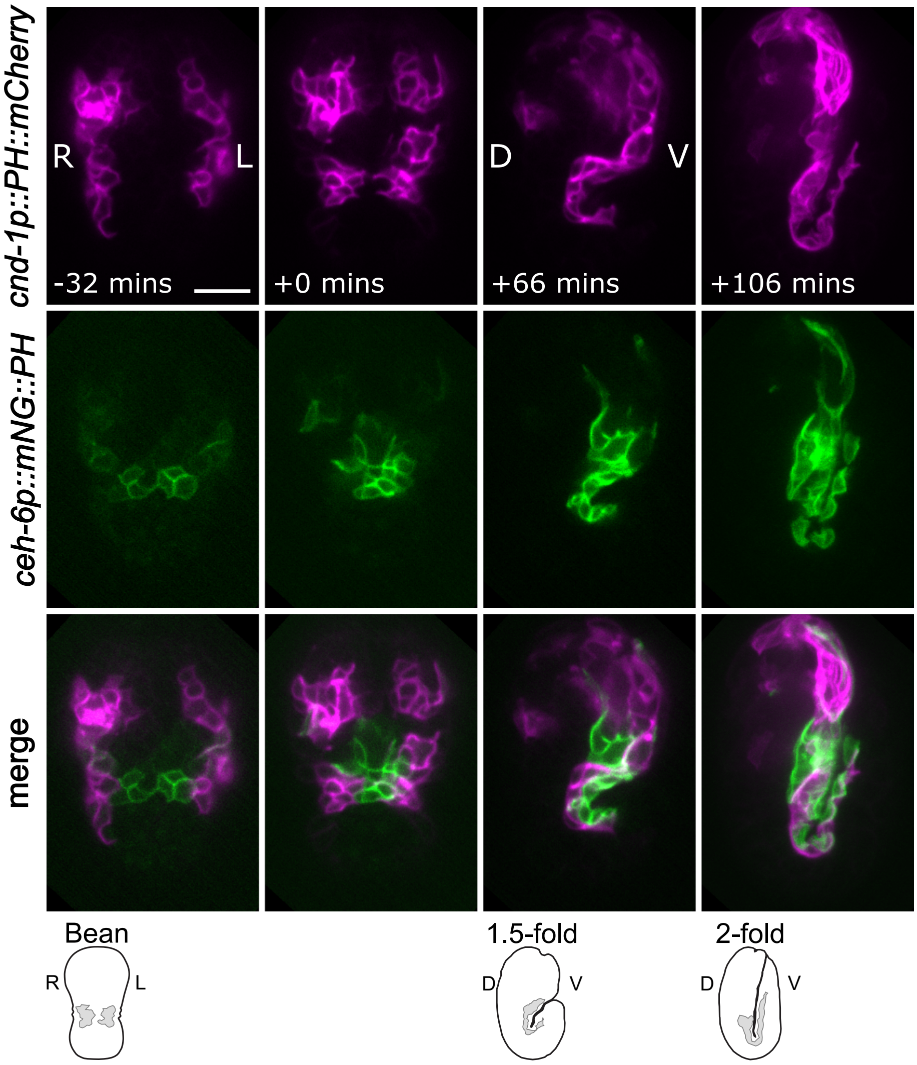

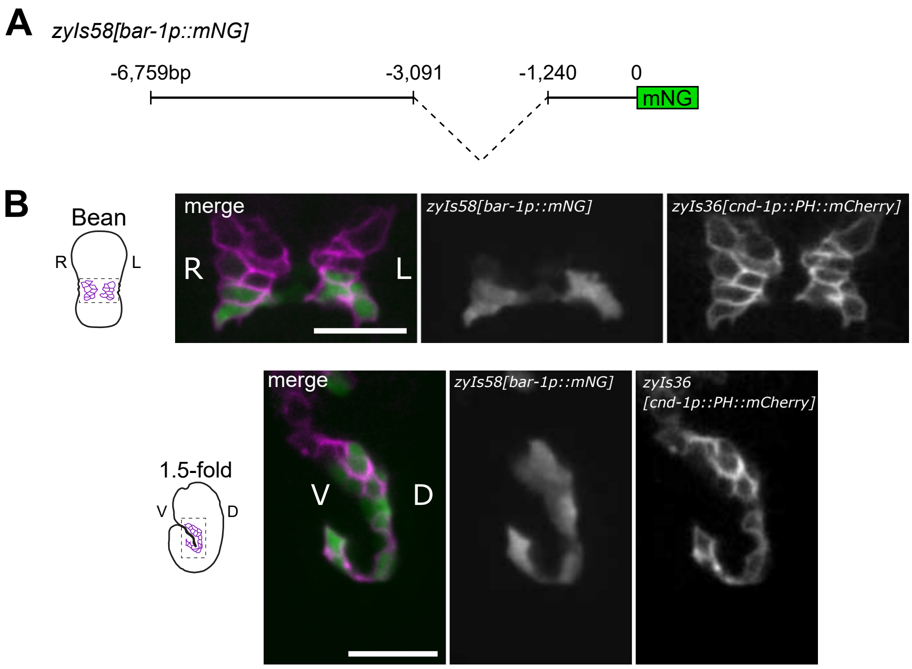
