## Supplemental Tables for "Notch-dependent left-right asymmetry in β-catenin-TCF activity instructs anteroposterior positioning of motor neurons in the *C. elegans* ventral nerve cord"

**Table S1.** Statistical analysis of DA, DB, and DD distribution at L1 in *C. elegans* strains analyzed in this study.

| Strain | Neuron | SD | Mean | Displacement * | Effect Size** | Significance *** |
|---|---|---|---|---|---|---|
| WT (n = 42) | DA1 | 1.749 | 8.63±0.55 | NA | NA | NA |
| | DA2 | 2.217 | 16.2±0.69 | NA | NA | NA |
| | DA3 | 2.564 | 26.99±0.8 | NA | NA | NA |
| | DA4 | 3.077 | 38.82±0.96 | NA | NA | NA |
| | DA5 | 3.16 | 53.22±0.98 | NA | NA | NA |
| | DA6 | 2.566 | 68.91±0.8 | NA | NA | NA |
| | DA7 | 1.497 | 82.97±0.47 | NA | NA | NA |
| | DA8 | 0.87 | 94.32±0.27 | NA | NA | NA |
| | DA9 | 0.721 | 95.09±0.22 | NA | NA | NA |
| | DD1 | 1.312 | 5.09±0.41 | NA | NA | NA |
| | DD2 | 2.616 | 22±0.82 | NA | NA | NA |
| | DD3 | 3.085 | 40.6±0.96 | NA | NA | NA |
| | DD4 | 3.099 | 60.86±0.97 | NA | NA | NA |
| | DD5 | 2.28 | 76.62±0.71 | NA | NA | NA |
| | DD6 | 1.095 | 93.02±0.34 | NA | NA | NA |
| | DB1 | 1.005 | 0.09±0.31 | NA | NA | NA |
| | DB2 | 1.332 | 5.62±0.42 | NA | NA | NA |
| | DB3 | 1.85 | 13.86±0.58 | NA | NA | NA |
| | DB4 | 2.819 | 30.1±0.88 | NA | NA | NA |
| | DB5 | 3.446 | 48.24±1.07 | NA | NA | NA |
| | DB6 | 2.717 | 66.15±0.85 | NA | NA | NA |
| | DB7 | 1.616 | 80.8±0.5 | NA | NA | NA |
| *bar-1(ga80)* (n = 45) | DA1 | 2.145 | 9.7±0.64 | 1.07 | 0.53 | p<0.05 |
| | DA2 | 3.556 | 15.45±1.07 | -0.76 | -0.25 | NS |
| | DA3 | 4.507 | 20.57±1.35 | -6.42 | -1.31 | p<0.001 |
| | DA4 | 2.832 | 39±0.85 | 0.18 | 0.06 | NS |
| | DA5 | 4.166 | 51.98±1.25 | -1.24 | -0.33 | NS |
| | DA6 | 2.613 | 68.73±0.79 | -0.18 | -0.07 | NS |
| | DA7 | 1.818 | 83.08±0.55 | 0.11 | 0.07 | NS |
| | DA8 | 0.592 | 94.41±0.18 | 0.1 | 0.13 | NS |
| | DA9 | 0.623 | 95.13±0.19 | 0.05 | 0.07 | NS |
| | DD1 | 1.715 | 5.39±0.52 | 0.3 | 0.19 | NS |
| | DD2 | 3.009 | 12.35±0.9 | -9.66 | -1.72 | p<0.001 |
| | DD3 | 3.047 | 40.82±0.92 | 0.22 | 0.07 | NS |
| | DD4 | 3.677 | 59.81±1.1 | -1.05 | -0.31 | NS |
| | DD5 | 2.553 | 75.55±0.77 | -1.07 | -0.43 | NS |
| | DD6 | 2.25 | 92.39±0.68 | -0.63 | -0.35 | NS |

| | | | | | | |
|---|---|---|---|---|---|---|
| | DB1 | 1.568 | 0.53±0.47 | 0.44 | 0.33 | NS |
| | DB2 | 1.676 | 6.88±0.5 | 1.26 | 0.77 | p<0.001 |
| | DB3 | 2.414 | 11.57±0.73 | -2.3 | -0.94 | p<0.001 |
| | DB4 | 2.662 | 30.29±0.8 | 0.19 | 0.07 | NS |
| | DB5 | 3.1 | 47.68±0.93 | -0.56 | -0.17 | NS |
| | DB6 | 2.666 | 65.8±0.8 | -0.35 | -0.13 | NS |
| | DB7 | 1.961 | 80.5±0.59 | -0.3 | -0.17 | NS |
| *bar-1; pop-1* (n = 43) | DA1 | 2.007 | 10.66±0.62 | 2.03 | 0.95 | p<0.001 |
| | DA2 | 2.652 | 16.87±0.82 | 0.67 | 0.27 | NS |
| | DA3 | 3.302 | 21.61±1.02 | -5.38 | -1.35 | p<0.001 |
| | DA4 | 4.199 | 37.24±1.29 | -1.58 | -0.42 | NS |
| | DA5 | 3.675 | 50.6±1.13 | -2.61 | -0.71 | p<0.01 |
| | DA6 | 3.053 | 67.03±0.94 | -1.88 | -0.64 | p<0.01 |
| | DA7 | 1.979 | 81.73±0.61 | -1.24 | -0.67 | p<0.01 |
| | DA8 | 0.769 | 93.72±0.24 | -0.6 | -0.69 | p<0.01 |
| | DA9 | 0.743 | 94.47±0.23 | -0.62 | -0.78 | p<0.001 |
| | DD1 | 1.655 | 6.29±0.51 | 1.2 | 0.75 | p<0.01 |
| | DD2 | 3.447 | 13.73±1.06 | -8.27 | -1.6 | p<0.001 |
| | DD3 | 3.069 | 39.71±0.94 | -0.89 | -0.29 | NS |
| | DD4 | 4.666 | 56.9±1.44 | -3.96 | -0.9 | p<0.001 |
| | DD5 | 2.601 | 74.6±0.8 | -2.03 | -0.77 | p<0.01 |
| | DD6 | 3.749 | 90.67±1.15 | -2.35 | -0.78 | p<0.001 |
| | DB1 | 1.795 | 1.59±0.55 | 1.5 | 0.92 | p<0.001 |
| | DB2 | 1.697 | 6.76±0.52 | 1.14 | 0.7 | p<0.01 |
| | DB3 | 2.348 | 11.49±0.72 | -2.37 | -0.98 | p<0.001 |
| | DB4 | 2.451 | 30.37±0.75 | 0.27 | 0.1 | NS |
| | DB5 | 2.494 | 46.62±0.77 | -1.62 | -0.53 | p<0.05 |
| | DB6 | 2.739 | 64.8±0.84 | -1.35 | -0.48 | NS |
| | DB7 | 1.857 | 78.99±0.57 | -1.81 | -0.92 | p<0.001 |
| *bar-1; pry-1(mu38)* (n = 39) | DA1 | 2.199 | 8.1±0.71 | -0.54 | -0.27 | NS |
| | DA2 | 2.663 | 15.83±0.86 | -0.37 | -0.15 | NS |
| | DA3 | 3.227 | 20.14±1.05 | -6.85 | -1.53 | p<0.001 |
| | DA4 | 3.024 | 38.24±0.98 | -0.58 | -0.19 | NS |
| | DA5 | 2.898 | 50.77±0.94 | -2.44 | -0.75 | p<0.01 |
| | DA6 | 2.369 | 67.86±0.77 | -1.05 | -0.42 | NS |
| | DA7 | 1.348 | 83.06±0.44 | 0.08 | 0.06 | NS |
| | DA8 | 0.664 | 94.06±0.22 | -0.26 | -0.33 | NS |
| | DA9 | 0.659 | 94.81±0.21 | -0.28 | -0.4 | NS |
| | DD1 | 1.627 | 5.46±0.53 | 0.37 | 0.25 | NS |
| | DD2 | 2.902 | 10.73±0.94 | -11.27 | -1.79 | p<0.001 |
| | DD3 | 4.396 | 40.09±1.43 | -0.51 | -0.14 | NS |
| | DD4 | 5.837 | 55.82±1.89 | -5.05 | -0.96 | p<0.001 |
| | DD5 | 1.641 | 74.78±0.53 | -1.85 | -0.84 | p<0.001 |

| | | | | | | |
|---|---|---|---|---|---|---|
| | DD6 | 2.318 | 91.83±0.75 | -1.19 | -0.63 | p<0.05 |
| | DB1 | 1.752 | 0.58±0.57 | 0.49 | 0.34 | NS |
| | DB2 | 1.441 | 6.12±0.47 | 0.5 | 0.35 | NS |
| | DB3 | 2.267 | 10.53±0.73 | -3.33 | -1.26 | p<0.001 |
| | DB4 | 4.034 | 30.33±1.31 | 0.23 | 0.07 | NS |
| | DB5 | 3.046 | 47.21±0.99 | -1.03 | -0.31 | NS |
| | DB6 | 2.302 | 65.1±0.75 | -1.06 | -0.41 | NS |
| | DB7 | 1.707 | 79.96±0.55 | -0.84 | -0.49 | NS |
| *bar-1; pry-1(zy103)* (n = 44) | DA1 | 1.947 | 9±0.59 | 0.36 | 0.2 | NS |
| | DA2 | 3.255 | 15.95±0.99 | -0.25 | -0.09 | NS |
| | DA3 | 3.458 | 20.54±1.05 | -6.45 | -1.45 | p<0.001 |
| | DA4 | 2.907 | 38.26±0.88 | -0.56 | -0.19 | NS |
| | DA5 | 4.136 | 51.95±1.26 | -1.26 | -0.34 | NS |
| | DA6 | 3.169 | 69.06±0.96 | 0.15 | 0.05 | NS |
| | DA7 | 1.512 | 83.53±0.46 | 0.55 | 0.36 | NS |
| | DA8 | 0.596 | 94.41±0.18 | 0.1 | 0.13 | NS |
| | DA9 | 0.585 | 95.15±0.18 | 0.07 | 0.1 | NS |
| | DD1 | 1.275 | 5.77±0.39 | 0.68 | 0.51 | p<0.05 |
| | DD2 | 2.654 | 12.32±0.81 | -9.68 | -1.75 | p<0.001 |
| | DD3 | 2.989 | 40.5±0.91 | -0.1 | -0.03 | NS |
| | DD4 | 5.938 | 58.56±1.81 | -2.3 | -0.47 | NS |
| | DD5 | 2.47 | 75.96±0.75 | -0.66 | -0.28 | NS |
| | DD6 | 2.384 | 92.29±0.72 | -0.73 | -0.38 | NS |
| | DB1 | 1.769 | 0.86±0.54 | 0.77 | 0.52 | p<0.05 |
| | DB2 | 1.168 | 6.38±0.36 | 0.76 | 0.59 | p<0.05 |
| | DB3 | 2.375 | 10.89±0.72 | -2.97 | -1.14 | p<0.001 |
| | DB4 | 3.237 | 30.57±0.98 | 0.47 | 0.15 | NS |
| | DB5 | 3.313 | 47.45±1.01 | -0.79 | -0.23 | NS |
| | DB6 | 2.931 | 65.72±0.89 | -0.43 | -0.15 | NS |
| | DB7 | 1.75 | 80.47±0.53 | -0.33 | -0.2 | NS |
| *mig-14(ga62)* (n = 42) | DA1 | 5.914 | 13.02±1.84 | 4.38 | 0.9 | p<0.001 |
| | DA2 | 5.443 | 21.28±1.7 | 5.08 | 1.05 | p<0.001 |
| | DA3 | 3.859 | 27.63±1.2 | 0.64 | 0.2 | NS |
| | DA4 | 5.098 | 39.79±1.59 | 0.97 | 0.23 | NS |
| | DA5 | 5.959 | 55.99±1.86 | 2.78 | 0.56 | p<0.05 |
| | DA6 | 5.989 | 70.45±1.87 | 1.54 | 0.33 | NS |
| | DA7 | 2.78 | 83.44±0.87 | 0.47 | 0.21 | NS |
| | DA8 | 1.125 | 93.63±0.35 | -0.69 | -0.65 | p<0.01 |
| | DA9 | 1.329 | 94.51±0.41 | -0.57 | -0.52 | p<0.05 |
| | DD1 | 2.178 | 6.01±0.68 | 0.91 | 0.49 | NS |
| | DD2 | 3.372 | 21.67±1.05 | -0.33 | -0.11 | NS |
| | DD3 | 4.561 | 39.6±1.42 | -1 | -0.26 | NS |
| | DD4 | 4.666 | 60.28±1.45 | -0.58 | -0.15 | NS |

| | | | | | | |
|---|---|---|---|---|---|---|
| | DD5 | 3.816 | 77.04±1.19 | 0.42 | 0.13 | NS |
| | DD6 | 2.526 | 92.06±0.79 | -0.96 | -0.48 | NS |
| | DB1 | 1.554 | -1.67±0.48 | -1.76 | -1.12 | p<0.001 |
| | DB2 | 2.701 | 8.24±0.84 | 2.63 | 1.05 | p<0.001 |
| | DB3 | 3.901 | 18.27±1.22 | 4.41 | 1.17 | p<0.001 |
| | DB4 | 3.648 | 31.58±1.14 | 1.48 | 0.45 | NS |
| | DB5 | 4.317 | 45.57±1.35 | -2.67 | -0.65 | p<0.01 |
| | DB6 | 3.751 | 65.72±1.17 | -0.43 | -0.13 | NS |
| | DB7 | 2.781 | 80.82±0.87 | 0.02 | 0.01 | NS |
| *mig-14; bar-1* (n = 42) | DA1 | 2.792 | 11.46±0.87 | 2.82 | 1.04 | p<0.001 |
| | DA2 | 3.434 | 23.2±1.07 | 6.99 | 1.54 | p<0.001 |
| | DA3 | 2.874 | 28.38±0.9 | 1.39 | 0.5 | NS |
| | DA4 | 3.042 | 38.49±0.95 | -0.33 | -0.11 | NS |
| | DA5 | 4.939 | 53.4±1.54 | 0.19 | 0.05 | NS |
| | DA6 | 3.809 | 68.55±1.19 | -0.36 | -0.11 | NS |
| | DA7 | 2.285 | 82.05±0.71 | -0.92 | -0.47 | NS |
| | DA8 | 0.855 | 93.36±0.27 | -0.96 | -0.98 | p<0.001 |
| | DA9 | 0.815 | 94.04±0.25 | -1.05 | -1.13 | p<0.001 |
| | DD1 | 3.004 | 7.49±0.94 | 2.4 | 0.92 | p<0.001 |
| | DD2 | 3.402 | 16.95±1.06 | -5.06 | -1.28 | p<0.001 |
| | DD3 | 3.772 | 38.69±1.18 | -1.91 | -0.54 | p<0.05 |
| | DD4 | 7.838 | 54.56±2.44 | -6.3 | -0.94 | p<0.001 |
| | DD5 | 2.986 | 74.87±0.93 | -1.75 | -0.63 | p<0.05 |
| | DD6 | 2.643 | 91.56±0.82 | -1.46 | -0.68 | p<0.01 |
| | DB1 | 2.792 | -0.79±0.87 | -0.88 | -0.41 | NS |
| | DB2 | 3.364 | 8.93±1.05 | 3.31 | 1.09 | p<0.001 |
| | DB3 | 4.103 | 17.11±1.28 | 3.25 | 0.91 | p<0.001 |
| | DB4 | 3.251 | 30.37±1.01 | 0.26 | 0.09 | NS |
| | DB5 | 3.247 | 44.78±1.01 | -3.46 | -0.92 | p<0.001 |
| | DB6 | 3.83 | 64.64±1.19 | -1.51 | -0.45 | NS |
| | DB7 | 2.477 | 79.21±0.77 | -1.59 | -0.71 | p<0.01 |
| *pop-1(hu9)* (n = 44) | DA1 | 1.506 | 9.03±0.46 | 0.39 | 0.24 | NS |
| | DA2 | 2.531 | 16.49±0.77 | 0.28 | 0.12 | NS |
| | DA3 | 2.848 | 23.1±0.87 | -3.89 | -1.17 | p<0.001 |
| | DA4 | 2.531 | 39.73±0.77 | 0.92 | 0.32 | NS |
| | DA5 | 3.14 | 53.21±0.95 | 0 | 0 | NS |
| | DA6 | 2.368 | 69.62±0.72 | 0.71 | 0.29 | NS |
| | DA7 | 1.731 | 83.65±0.53 | 0.68 | 0.41 | NS |
| | DA8 | 0.867 | 93.86±0.26 | -0.45 | -0.51 | p<0.05 |
| | DA9 | 0.896 | 94.57±0.27 | -0.51 | -0.6 | p<0.05 |
| | DD1 | 1.155 | 5.09±0.35 | -0.01 | -0.01 | NS |
| | DD2 | 2.493 | 17.29±0.76 | -4.71 | -1.36 | p<0.001 |
| | DD3 | 2.824 | 41.12±0.86 | 0.52 | 0.18 | NS |

| | | | | | | |
|---|---|---|---|---|---|---|
| | DD4 | 2.761 | 60.93±0.84 | 0.07 | 0.02 | NS |
| | DD5 | 2.507 | 76.4±0.76 | -0.23 | -0.1 | NS |
| | DD6 | 1.619 | 92.51±0.49 | -0.5 | -0.36 | NS |
| | DB1 | 1.411 | 0.24±0.43 | 0.15 | 0.12 | NS |
| | DB2 | 1.519 | 5.74±0.46 | 0.12 | 0.08 | NS |
| | DB3 | 2.352 | 13.24±0.71 | -0.62 | -0.29 | NS |
| | DB4 | 2.37 | 30.53±0.72 | 0.43 | 0.16 | NS |
| | DB5 | 3.023 | 47.9±0.92 | -0.34 | -0.1 | NS |
| | DB6 | 2.509 | 66.73±0.76 | 0.58 | 0.22 | NS |
| | DB7 | 2.016 | 80.89±0.61 | 0.09 | 0.05 | NS |
| | DA1 | 2.496 | 10.95±0.77 | 2.31 | 0.95 | p<0.001 |
| | DA2 | 4.175 | 23.3±1.28 | 7.1 | 1.45 | p<0.001 |
| | DA3 | 2.673 | 29.19±0.82 | 2.2 | 0.78 | p<0.001 |
| | DA4 | 3.488 | 41.18±1.07 | 2.36 | 0.68 | p<0.01 |
| | DA5 | 3.467 | 53±1.07 | -0.22 | -0.07 | NS |
| | DA6 | 2.156 | 68.9±0.66 | -0.01 | 0 | NS |
| | DA7 | 1.656 | 82.53±0.51 | -0.44 | -0.28 | NS |
| | DA8 | 0.541 | 93.63±0.17 | -0.68 | -0.86 | p<0.001 |
| | DA9 | 0.576 | 94.4±0.18 | -0.69 | -0.94 | p<0.001 |
| | DD1 | 5.38 | 11.44±1.66 | 6.35 | 1.26 | p<0.001 |
| *pry-1(mu38)* (n = 43) | DD2 | 2.734 | 22.53±0.84 | 0.52 | 0.2 | NS |
| | DD3 | 3.558 | 41.69±1.1 | 1.09 | 0.32 | NS |
| | DD4 | 3.092 | 60.74±0.95 | -0.12 | -0.04 | NS |
| | DD5 | 1.576 | 76.72±0.49 | 0.1 | 0.05 | NS |
| | DD6 | 1.518 | 92.07±0.47 | -0.95 | -0.68 | p<0.01 |
| | DB1 | 1.442 | 0.84±0.44 | 0.75 | 0.58 | p<0.05 |
| | DB2 | 1.702 | 6.69±0.52 | 1.07 | 0.66 | p<0.01 |
| | DB3 | 2.271 | 16.24±0.7 | 2.37 | 1 | p<0.001 |
| | DB4 | 2.51 | 31.52±0.77 | 1.42 | 0.52 | p<0.05 |
| | DB5 | 3.165 | 47.43±0.97 | -0.81 | -0.25 | NS |
| | DB6 | 2.768 | 66.33±0.85 | 0.18 | 0.07 | NS |
| | DB7 | 1.749 | 80.43±0.54 | -0.37 | -0.22 | NS |
| | DA1 | 2.362 | 10.65±0.64 | 2.02 | 0.87 | p<0.001 |
| | DA2 | 3.184 | 20.83±0.87 | 4.62 | 1.28 | p<0.001 |
| | DA3 | 2.592 | 28.39±0.71 | 1.4 | 0.53 | p<0.05 |
| | DA4 | 3.131 | 39.7±0.85 | 0.88 | 0.28 | NS |
| | DA5 | 3.55 | 53.23±0.97 | 0.01 | 0 | NS |
| *pry-1(zy103)* (n = 54) | DA6 | 3.349 | 68.83±0.91 | -0.09 | -0.03 | NS |
| | DA7 | 2.139 | 82.88±0.58 | -0.09 | -0.05 | NS |
| | DA8 | 0.752 | 94.54±0.21 | 0.22 | 0.28 | NS |
| | DA9 | 0.705 | 95.37±0.19 | 0.28 | 0.39 | NS |
| | DD1 | 3.241 | 8.41±0.88 | 3.31 | 1.08 | p<0.001 |
| | DD2 | 2.192 | 23.41±0.6 | 1.41 | 0.57 | p<0.05 |

| | | | | | | |
|---|---|---|---|---|---|---|
| | DD3 | 3.128 | 41.48±0.85 | 0.88 | 0.28 | NS |
| | DD4 | 3.664 | 60.43±1 | -0.43 | -0.13 | NS |
| | DD5 | 2.576 | 76.72±0.7 | 0.1 | 0.04 | NS |
| | DD6 | 0.982 | 93.35±0.27 | 0.34 | 0.32 | NS |
| | DB1 | 1.635 | 0.1±0.45 | 0.01 | 0 | NS |
| | DB2 | 1.488 | 6.62±0.41 | 1 | 0.67 | p<0.01 |
| | DB3 | 1.922 | 16.59±0.52 | 2.73 | 1.18 | p<0.001 |
| | DB4 | 2.821 | 31.54±0.77 | 1.43 | 0.5 | p<0.05 |
| | DB5 | 3.173 | 48.01±0.87 | -0.23 | -0.07 | NS |
| | DB6 | 3.307 | 65.78±0.9 | -0.38 | -0.12 | NS |
| | DB7 | 2.364 | 80.53±0.65 | -0.27 | -0.13 | NS |
| *pry-1(zy103); pop-1* (n = 42) | DA1 | 1.814 | 8.94±0.57 | 0.31 | 0.17 | NS |
| | DA2 | 3.186 | 18.48±0.99 | 2.28 | 0.77 | p<0.01 |
| | DA3 | 3.614 | 23.74±1.13 | -3.25 | -0.92 | p<0.001 |
| | DA4 | 3.279 | 38.17±1.02 | -0.64 | -0.2 | NS |
| | DA5 | 2.999 | 52.14±0.93 | -1.08 | -0.35 | NS |
| | DA6 | 2.315 | 68.22±0.72 | -0.69 | -0.28 | NS |
| | DA7 | 1.404 | 82.86±0.44 | -0.11 | -0.08 | NS |
| | DA8 | 0.595 | 93.88±0.19 | -0.44 | -0.57 | p<0.05 |
| | DA9 | 0.571 | 94.65±0.18 | -0.44 | -0.64 | p<0.01 |
| | DD1 | 1.99 | 6.18±0.62 | 1.08 | 0.61 | p<0.05 |
| | DD2 | 2.839 | 16.94±0.88 | -5.06 | -1.36 | p<0.001 |
| | DD3 | 4.176 | 39.57±1.3 | -1.03 | -0.28 | NS |
| | DD4 | 3.996 | 58.73±1.25 | -2.13 | -0.57 | p<0.05 |
| | DD5 | 1.885 | 75.17±0.59 | -1.46 | -0.66 | p<0.01 |
| | DD6 | 2.381 | 91.63±0.74 | -1.39 | -0.71 | p<0.01 |
| | DB1 | 1.568 | 0.78±0.49 | 0.69 | 0.51 | p<0.05 |
| | DB2 | 1.911 | 5.64±0.6 | 0.02 | 0.01 | NS |
| | DB3 | 2.342 | 14.64±0.73 | 0.78 | 0.37 | NS |
| | DB4 | 3.145 | 29.45±0.98 | -0.65 | -0.22 | NS |
| | DB5 | 2.944 | 47.18±0.92 | -1.06 | -0.33 | NS |
| | DB6 | 2.734 | 65.43±0.85 | -0.72 | -0.26 | NS |
| | DB7 | 1.713 | 80.17±0.53 | -0.63 | -0.37 | NS |
| *zyIs59* (n = 47) | DA1 | 1.329 | 8.94±0.39 | 0.31 | 0.2 | NS |
| | DA2 | 1.556 | 17.32±0.46 | 1.11 | 0.57 | p<0.05 |
| | DA3 | 2.065 | 27.05±0.61 | 0.06 | 0.03 | NS |
| | DA4 | 2.044 | 38.93±0.6 | 0.11 | 0.04 | NS |
| | DA5 | 2.227 | 52.46±0.65 | -0.76 | -0.28 | NS |
| | DA6 | 2.131 | 68.21±0.63 | -0.7 | -0.3 | NS |
| | DA7 | 1.45 | 82.37±0.43 | -0.61 | -0.4 | NS |
| | DA8 | 0.668 | 94.31±0.2 | 0 | 0 | NS |
| | DA9 | 0.664 | 94.93±0.19 | -0.16 | -0.22 | NS |
| | DD1 | 1.166 | 5.36±0.34 | 0.26 | 0.21 | NS |

| | | | | | | |
|---|---|---|---|---|---|---|
| | DD2 | 1.714 | 22.98±0.5 | 0.98 | 0.44 | NS |
| | DD3 | 2.497 | 41.34±0.73 | 0.73 | 0.26 | NS |
| | DD4 | 3.282 | 61.4±0.96 | 0.54 | 0.17 | NS |
| | DD5 | 1.829 | 76.56±0.54 | -0.07 | -0.03 | NS |
| | DD6 | 2.38 | 92.33±0.7 | -0.69 | -0.36 | NS |
| | DB1 | 1.274 | -0.06±0.37 | -0.15 | -0.13 | NS |
| | DB2 | 1.36 | 6.09±0.4 | 0.47 | 0.34 | NS |
| | DB3 | 1.421 | 14.64±0.42 | 0.77 | 0.46 | NS |
| | DB4 | 1.959 | 30.32±0.58 | 0.21 | 0.09 | NS |
| | DB5 | 2.015 | 47.17±0.59 | -1.07 | -0.38 | NS |
| | DB6 | 2.376 | 64.88±0.7 | -1.27 | -0.49 | p<0.05 |
| | DB7 | 1.645 | 80.1±0.48 | -0.7 | -0.42 | NS |
| *zyIs59; pry-1(zy103)* (n = 48) | DA1 | 2.619 | 12.48±0.76 | 3.85 | 1.3 | p<0.001 |
| | DA2 | 4.004 | 22.58±1.16 | 6.38 | 1.39 | p<0.001 |
| | DA3 | 3.812 | 30.26±1.11 | 3.27 | 0.89 | p<0.001 |
| | DA4 | 4.413 | 43±1.28 | 4.18 | 0.96 | p<0.001 |
| | DA5 | 4.089 | 54.87±1.19 | 1.66 | 0.44 | NS |
| | DA6 | 3.198 | 69.81±0.93 | 0.9 | 0.31 | NS |
| | DA7 | 2.425 | 83.16±0.7 | 0.19 | 0.09 | NS |
| | DA8 | 0.64 | 94.9±0.19 | 0.59 | 0.73 | p<0.01 |
| | DA9 | 0.672 | 95.56±0.2 | 0.48 | 0.65 | p<0.01 |
| | DD1 | 4.821 | 12.83±1.4 | 7.74 | 1.46 | p<0.001 |
| | DD2 | 3.095 | 25.69±0.9 | 3.68 | 1.08 | p<0.001 |
| | DD3 | 7.634 | 53.26±2.22 | 12.65 | 1.46 | p<0.001 |
| | DD4 | 5.458 | 72.76±1.58 | 11.9 | 1.59 | p<0.001 |
| | DD5 | 3.375 | 84.26±0.98 | 7.64 | 1.59 | p<0.001 |
| | DD6 | 1.109 | 93.7±0.32 | 0.68 | 0.6 | p<0.05 |
| | DB1 | 1.806 | 1.4±0.52 | 1.31 | 0.81 | p<0.001 |
| | DB2 | 2.15 | 7.5±0.62 | 1.88 | 0.92 | p<0.001 |
| | DB3 | 2.829 | 16.19±0.82 | 2.32 | 0.87 | p<0.001 |
| | DB4 | 4.024 | 33.31±1.17 | 3.21 | 0.83 | p<0.001 |
| | DB5 | 4.345 | 50.26±1.26 | 2.02 | 0.5 | p<0.05 |
| | DB6 | 3.279 | 66.25±0.95 | 0.09 | 0.03 | NS |
| | DB7 | 2.625 | 80.36±0.76 | -0.44 | -0.2 | NS |

*Displacement is calculated by subtracting the mean mutant position with the mean WT position of the corresponding neuron. Positive and negative displacements correspond to posterior and anterior position shifts, respectively.

** Cohen d’s Effect Size

*** Welch’s t-test with Benjamini-Hochberg (FDR) adjustment applied across all tests in this table

**Table S2.** Oligonucleotides used to generate sgRNA and HDR constructs for CRISPR/Cas9 genome editing using SEC-based and co-CRISPR approaches.

| Oligo | DNA Oligo Sequence (5' - 3') | Use |
|---|---|---|
| W1 | TCAGAAAACTGGGATGGACCGTTTTAGAGCTAGAAATAGCAAGT | sgRNA to target N-terminus of *bar-1* for *GFP::bar-1* knock-in |
| W2 | ACGTTGTAAAACGACGGCCAGTCGCCGGCATGTGTGCCTTCTCGTTTCAC | HDR repair plasmid for *GFP::bar-1* knock-in |
| W3 | TCCAGTGAACAATTCTTCTCCTTTACTCATCCCAGTTTTCTGAAAAAAAAAGCCA | HDR repair plasmid for *GFP::bar-1* knock-in |
| W4 | CGTGATTACAAGGATGACGATGACAAGAGAATGGACCTGGATCCGAACC | HDR repair plasmid for *GFP::bar-1* knock-in |
| W5 | TCACACAGGAAACAGCTATGACCATGTTATTGTTATACTGCCCATCCGCT | HDR repair plasmid for *GFP::bar-1* knock-in |
| W6 | GGAGCATCGATGGTCTCCAAGTTTAAGAGCTATGCTGGAA | sgRNA to target mNG in *pry-1(cp383)* |
| W7 | TCGTGGTGGGCCTCTGACACATGCAGCTC | mNG::T2A:: mScarlet-i::H2B::pry-1-3'UTR repair fragment for insertion into *cp383* |
| W8 | AATCCCACTACTTGCTGGAAGTGTACTTGG | mNG::T2A:: mScarlet-i::H2B::pry-1-3'UTR repair fragment for insertion into *cp383* |
| W9 | TTCCAGCAAGTAGTGGGATTATCGATTTTTGAAAAC | mNG::T2A:: mScarlet-i::H2B::pry-1-3'UTR repair fragment for insertion into *cp383* |
| W10 | GTGTCAGAGGCCCACCACGAAAACCTAC | mNG::T2A:: mScarlet-i::H2B::pry-1-3'UTR repair fragment for insertion into *cp383* |
| W11 | ACTGTTGGATGCCTGTGTAGGTTTAAGAGCTATGCTGGAA | sgRNA to target *cxTi10882* to generate *zySi6[bar-1p::mNG::PH]* |
| W12 | AAGTGAGTTTGCTACCATCAGTTTAAGAGCTATGCTGGAA | sgRNA to target *ttTi5605* to generate *zySi10[unc-30p::mNG::H2B]* |

**Table S3.** Mutants identified in a genetic screen for DD neuron position defects.

| Mutant allele | Chr | Position (WS285) | Mutation | Type | Effect |
|---|---|---|---|---|---|
| *bar-1(zy99)* | X | 7170307 | C to T | splicing | exon 2 donor site |
| *pry-1(zy103)* | I | 14175137 | G to A | nonsense | R 355 stop |
| *lin-12(zy127)* | III | 9064316 | A to T | missense | C 491 S |
| *lag-1(zy129)* | IV | 6108439 | G to A | missense | E 438 K (isoform A) |

**Table S4**. Statistics for *bar-1(ga80)*, *pry-1(zy103),* and *pry-1(mu38)* genetic interaction analyses

| **Compared to:** | | **DA1** | **DA2** | **DA3** | **DB1** | **DB2** | **DB3** | **DD1** | **DD2** | **DD3** |
|---|---|---|---|---|---|---|---|---|---|---|
| *bar-1(ga80); pry-1(mu38)* | *bar-1(ga80)* | * | ns | ns | | ns | ns | ns | ns | |
| *bar-1(ga80); pry-1(zy103)* | *bar-1(ga80)* | ns | ns | ns | | ns | ns | ns | ns | |
| *pry-1(mu38)* | *bar-1(ga80)* | ns | *** | *** | | ns | *** | *** | *** | |
| *pry-1(zy103)* | *bar-1(ga80)* | ns | *** | *** | | ns | *** | *** | *** | |
| *bar-1(ga80); pry-1(zy103)* | *bar-1(ga80); pry-1(mu38)* | ns | ns | ns | | ns | ns | ns | ns | |
| *pry-1(mu38)* | *bar-1(ga80); pry-1(mu38)* | *** | *** | *** | | ns | *** | *** | *** | |
| *pry-1(zy103)* | *bar-1(ga80); pry-1(mu38)* | *** | *** | *** | | ns | *** | *** | *** | |
| *pry-1(mu38)* | *bar-1(ga80); pry-1(zy103)* | *** | *** | *** | | ns | *** | *** | *** | |
| *pry-1(zy103)* | *bar-1(ga80); pry-1(zy103)* | ** | *** | *** | | ns | *** | *** | *** | |
| *pry-1(zy103)* | *pry-1(mu38)* | ns | ** | ns | | ns | ns | *** | ns | |
| **ANOVA** | | *** | *** | *** | ns | ** | *** | *** | *** | ns |

Comparisons against WT are omitted. Statistics used was an ANOVA with Tukey's HSD post-hoc test. p<0.05 (*), p<0.01(**), p<0.001(***).

**Table S5**. Statistics for *pry-1(zy103)* and *pop-1(hu9)* genetic interaction analyses

| **Compared to:** | | **DA1** | **DA2** | **DA3** | **DB1** | **DB2** | **DB3** | **DD1** | **DD2** |
|---|---|---|---|---|---|---|---|---|---|
| *pry-1* | *pop-1* | *** | *** | *** | | * | *** | *** | *** |
| *pop-1; pry-1* | *pop-1* | ns | ** | ns | | ns | * | ns | ns |
| *pop-1; pry-1* | *pry-1* | *** | *** | *** | | * | *** | *** | *** |
| **ANOVA** | | *** | *** | *** | ns | ** | *** | *** | *** |

Comparisons against WT are omitted. Statistics used was an ANOVA with Tukey's HSD post-hoc test. p<0.05 (*), p<0.01(**), p<0.001(***).

**Table S6**. Statistics for *bar-1(ga80)* and *pop-1(hu9)* genetic interaction analyses

| **Compared to:** | | **DA1** | **DA2** | **DA3** | **DB1** | **DB2** | **DB3** | **DD1** | **DD2** |
|---|---|---|---|---|---|---|---|---|---|
| *pop-1; bar-1* | *bar-1* | ns | | ns | ** | ns | ns | * | ns |
| *pop-1* | *bar-1* | ns | | ** | ns | ** | ** | ns | *** |
| *pop-1* | *pop-1; bar-1* | *** | | ns | *** | * | ** | ** | *** |
| **ANOVA** | | *** | ns | *** | *** | *** | *** | *** | *** |

Comparisons against WT are omitted. Statistics used was an ANOVA with Tukey's HSD post-hoc test. p<0.05 (*), p<0.01(**), p<0.001(***).

**Table S7**. Statistics for *bar-1(ga80)* and *mig-14(ga62)* genetic interaction analyses

| **Compared to:** | | **DA1** | **DA2** | **DA3** | **DB1** | **DB2** | **DB3** | **DD1** | **DD2** |
|---|---|---|---|---|---|---|---|---|---|
| *mig-14* | *bar-1* | *** | *** | *** | *** | * | *** | ns | *** |
| *mig-14; bar-1* | *bar-1* | ns | *** | *** | ** | *** | *** | *** | *** |
| *mig-14; bar-1* | *mig-14* | ns | ns | ns | ns | ns | ns | ** | *** |
| **ANOVA** | | *** | *** | *** | *** | *** | *** | *** | *** |

Comparisons against WT are omitted. Statistics used was an ANOVA with Tukey's HSD post-hoc test. p<0.05 (*), p<0.01(**), p<0.001(***).
